# Activatable prodrug for controlled release of an antimicrobial peptide via the proteases overexpressed in *Candida albicans* and *Porphyromonas gingivalis*

**DOI:** 10.1101/2023.11.27.568833

**Authors:** Lubna Amer, Maurice Retout, Jesse V. Jokerst

**Author notes:** **Corresponding author’s email:** (J.V.J.).

## Abstract

We report the controlled release of an antimicrobial peptide using enzyme-activatable prodrugs to treat and detect *Candida albicans* and *Porphyromonas gingivalis*. Our motivation lies in the prevalence of these microorganisms in the subgingival area where the frequency of fungal colonization increases with periodontal disease. This work is based on an antimicrobial peptide that is both therapeutic and induces a color change in a nanoparticle reporter. This antimicrobial peptide was then built into a zwitterionic prodrug that quenches its activity until activation by a protease inherent to these pathogens of interest: SAP9 or RgpB for *C. albicans* and *P. gingivalis*, respectively. We first confirmed that the intact zwitterionic prodrug has negligible toxicity to fungal, bacterial, and mammalian cells absent a protease trigger. Next, the therapeutic impact was assessed via disk diffusion and viability assays and showed a minimum inhibitory concentration of 3.1 – 16 µg/mL, which is comparable to the antimicrobial peptide alone (absent integration into prodrug). Finally, the zwitterionic design was exploited for colorimetric detection of *C. albicans* and *P. gingivalis* proteases. When the prodrugs were cleaved, the plasmonic nanoparticles aggregated causing a color change with a limit of detection of 10 nM with gold nanoparticles and 3 nM with silver nanoparticles. This approach has value as a convenient and selective protease sensing and protease-induced treatment mechanism based on bioinspired antimicrobial peptides.

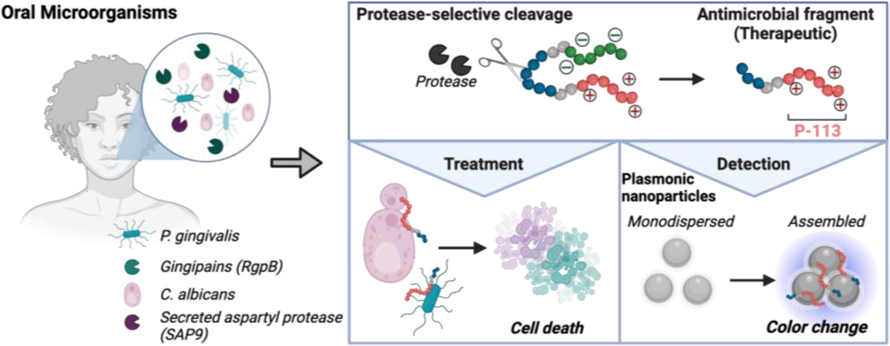

## INTRODUCTION

The oral cavity contains the highest diversity of microorganisms among all human microbial environments and serves as a primary site of colonization and cross-kingdom interactions between fungi and bacteria.[1, 2] The *Candida* genus is notorious for its ability to form biofilms and resist both host-derived and external antimicrobial molecules. *Candida* interacts with other oral microbes to promote hyphal formation and enhance the pathogenesis of both microorganisms.[3–8] Amongst the *Candida* genus, *Candida albicans* is the most commonly isolated species from the oral cavity and is responsible for most superficial and systemic fungal infections.[9, 10] *C. albicans* is an opportunistic yeast-like fungus that colonizes human mucosal surfaces often becoming pathogenic in immunocompromised patients. This can lead to recurrent or even disseminated infections with high mortality rates.[4] This pathogenic process is rooted in the colonization activity wherein *C. albicans* exploits an ability to switch morphology from yeast to hyphal form. Importantly, the hyphal form leads to a structured microbial biofilm that increases resistance to antimicrobial therapy.[3, 11–14] Biofilms that extend below the gingival margin become subgingival plaque.[15, 16] Here, pathogenic, gram-negative anaerobic bacteria such as *Porphyromonas gingivalis* can co-colonize the sulcus.[15]

*P. gingivalis* is a keystone bacteria in periodontal disease and involves an inflammatory process with various degrees of severity ranging from readily curable gingivitis to irreversible periodontitis.[15, 17–19] Data suggest that interactions between *C. albicans* and bacteria may modulate the clinical course of infection and negatively impact treatment.[20] Likewise, the rate of colonization of *C. albicans* at subgingival sites is significantly higher in subjects with chronic periodontitis; the rate of candidiasis increases with higher isolation frequencies of *P. gingivalis.*[21–23] However, treatment of *C. albicans* and *P. gingivalis* is quite different, and co-treatment has not yet been investigated in molecular detail.[21] Moreover, drug-resistant fungi are a growing public health threat with a mortality rate as high as 30% in *Candida* family (*C. auris*); thus, improved therapies are clearly needed.[9, 24] Likewise, treatment for *P. gingivalis*-based gingivitis is largely limited to lifestyle changes; thus, gingivitis relapses are common in susceptible patients leading to tooth loss and worse quality of life.[25]

Here, we report an activatable protease-responsive antimicrobial peptide as a novel therapeutic approach to treat and detect *C. albicans* and *P. gingivalis*. Antimicrobial peptides (**AMP**) are cationic, hydrophobic, and amphiphilic molecules and promising therapies against multi-drug resistant microorganisms. AMPs disrupt the cell membrane through electrostatic interactions with negatively charged components such as phospholipids and ergosterol.[26, 27] However, AMPs can be non-specific and damage host cells.[28, 29] Thus, we hypothesized that coating-activatable systems that are readily dependent on the pathogenesis of the microorganism, would overcome this limitation by retaining the pharmacokinetic properties through selectivity of treatment. This can be achieved through cleavable peptide substrates designed to release the AMP under direct exposure to a pathogenic protease.

More specifically, we tuned the prodrug for *C. albicans* to be responsive to secreted aspartic proteases (**SAP**), which govern the transition from harmless commensal to disease-causing pathogen.[22, 30–33] Similarly, gingipains (**gp**) are implicated in the pathogeneses of gingivitis and represent more than 85% of the total proteolytic activity of *P. gingivalis*.[34] Therefore, along with serving as a bacteria-presence signature, they are also relevant biomarkers of disease progression and development. We thus tuned a second prodrug to respond to gingipains. Finally, we designed the zwitterionic peptides to concurrently produce a point-of-care colorimetric signal that can report the presence of *C. albicans* or *P. gingivalis*. This color change is based on precious metal nanoparticles and plasmonic coupling leading to a change in optical behavior in the visible region.[34–37] In the work below, we characterize the prodrugs, their toxicity profiles, their selectivity to fungi and bacteria *in vitro*, and finally show the diagnostic performance of the sensor.

## RESULTS AND DISCUSSION

### Design of prodrugs specific to SAP9 or RgpB

Two activable prodrugs unique to *C. albicans* or *P. gingivalis* were designed comprising of three functional parts (**Figure 1A**). The first part (**Scheme 1** red) is the antimicrobial peptide (**AMP**; P-113; **AKRHHGYKRKFH; Table 1**). This 12-amino-acid derivative of Histatin-5 has antifungal activity comparable to that of the parent molecule with fewer sites prone to proteolytic degradation.[38] P-113’s electrostatic interference (net charge: +5) causes pore-like breakdowns of the cellular wall, leakage of intracellular contents, and ultimately, cell death (**Scheme 1**).[39] This process is rapid and broad spectrum[40, 41] and we hypothesized that it would have activity against *P. gingivalis* as well for joint antifungal and antibacterial treatment.[38, 42]

**Scheme 1.**
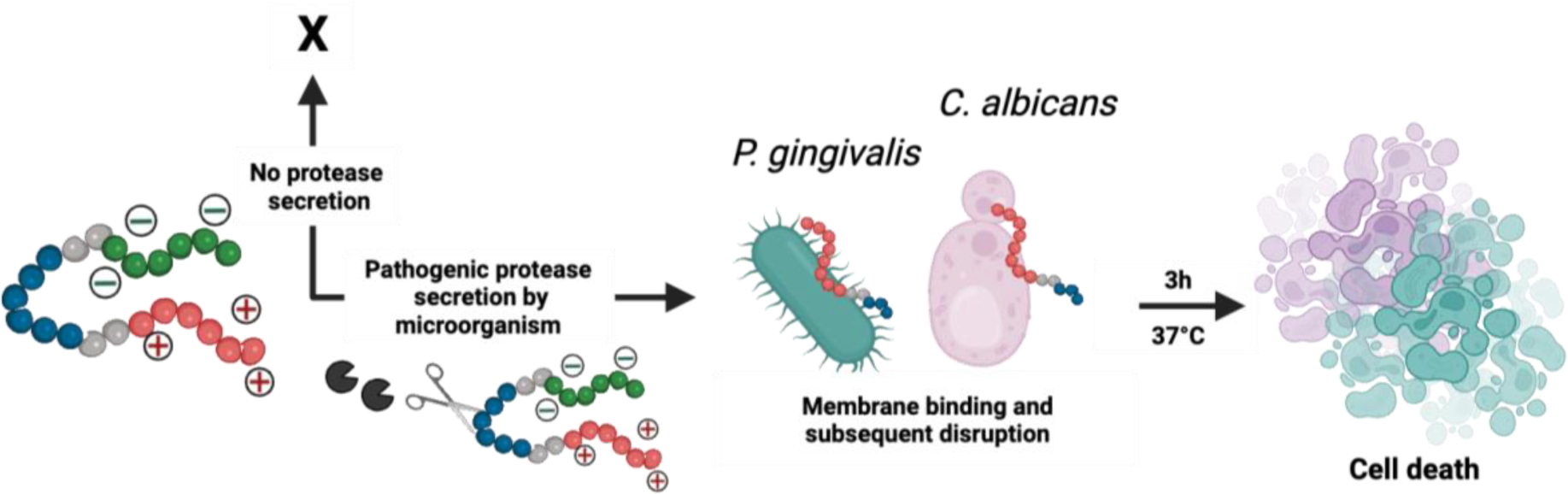
Descriptive schematic of activatable therapy. The three key components of the prodrug-peptide: a charge-shielding toxicity-quencher (DDDDDD) (green), a recognition sequence for SAP9 (<u>VKKK/DVVD</u>) or RgpB (<u>AGPR/ID</u>) (blue), and a therapy (P-113; **AKRHHGYKRKFH**) (red). The net charge of the intact prodrug and its therapy is 0 and +5, respectively. An expression of pathogenic proteases induces the cleavage of the prodrug, thus releasing the antimicrobial fragment from its toxicity quencher. In turn, the increased, positively charged antimicrobial fragment binds to the negatively charged components of the *C. albicans* or *P. gingivalis* cellular membrane thus causing interference in electrostatic structure and ultimately, cell death. Alternatively, under healthy conditions, the protease biomarkers are not expressed, the zwitterionic prodrug-peptide remains intact, and the antimicrobial activity does not activate.

**Figure 1.**
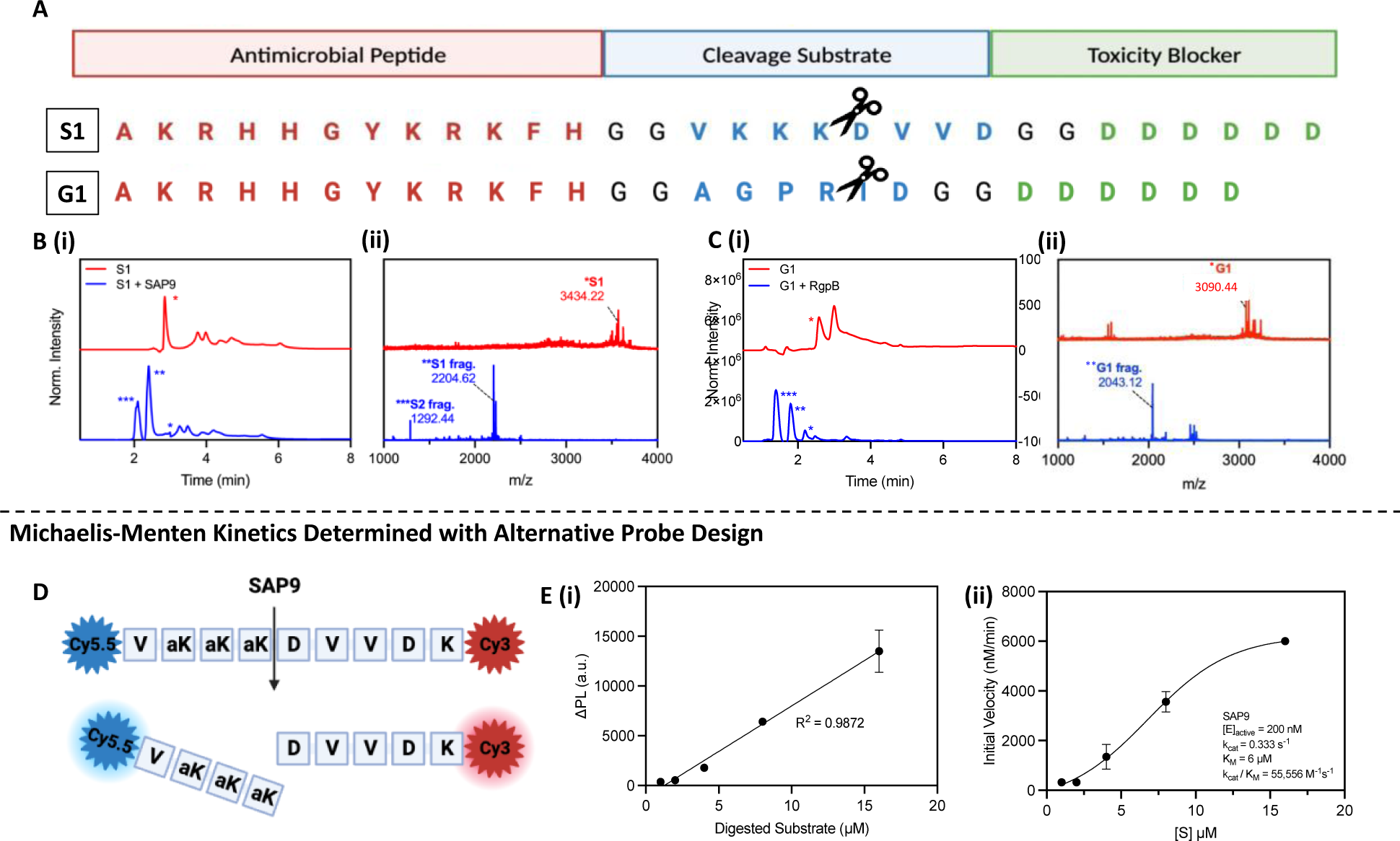
Activatable antimicrobial peptide design and cleavage characterization. **(A)** *C. albicans* and *P. gingivalis* cleavable prodrugs, S1 and G1, respectively (top to bottom) demonstrating the three main components: **(1)** A positively charged antimicrobial peptide, P-113 (red), **(2)** a cleavable peptide substrate (blue) and **(3)** a negatively charged toxicity quencher (green). Enzymes expressed by *C. albicans* (SAP9) can cleave the S1 prodrug after the third lysine in the cleavage substrate, thus resulting in the release of the antimicrobial fragment and subsequent therapeutic activity. Enzymes expressed by *P. gingivalis* (RgpB) can cleave the G1 prodrug after the arginine residue in the corresponding cleavage substrate and render the aforementioned effect. **(B) (i)** HPLC (220 nm wavelength monitoring) and **(ii)** MALDI-TOF spectra of S1 prodrug intact (red) and cleaved (blue, i.e., after 3 hours of incubation with 300 nM of SAP9) show a divide in the spectra with corresponding peaks at fragment mass. * S1 = S1 intact, ** S1 = S1 antimicrobial fragment, *** S1 = S1 toxicity blocker fragment. **(C) (i)** HPLC (220 nm wavelength monitoring) and **(ii)** MALDI-TOF spectra of G1 prodrug intact (red) and cleaved (blue, i.e., after 3 hours of incubation with 200 nM of RgpB) showing a divide in the spectra with corresponding peaks at fragment mass. * G1 = G1 intact, ** G1 = G1 antimicrobial fragment, *** G1 = G1 toxicity blocker fragment. **(D)** Structure of SAP9 cleavable heterodimer FRET probe with NHS-amine reacted cyanine dyes conjugated on either terminal to create strong fluorescence quenching when intact for estimates of Michalis-Menten parameters. **(E) (i)** Standard curve of linear increase in change in fluorescence signal as a result of enzyme incubation over time with respect to substrate concentration. **(ii)** Michaelis-Menten master curve describing the rate of reaction with an increase in substrate concentration to determine k_cat_/ K_M_, the second-order rate constant reaction rate of the enzyme-substrate complex to product. Here, a higher ratio suggests a higher rate of conversion. Inset parameters present: [E]_active_, the active enzyme concentration used, 200 nM. Catalytic constant, k_cat_, the number of probe molecules converted by enzyme per second. The Michaelis-Menten constant, K_M_, an inverse measurement of affinity.

**Table 1.** Key peptides. Peptide P-113 is an antimicrobial peptide that serves as a positive control. It is integrated into activatable peptides S1 and G1.

| Name | Sequence | Mass (Da) | Charge |
| --- | --- | --- | --- |
| P-113 | Ac-AKRHHGYKRKFH | 1604.8891 | +5 |
| *C. albicans*<br>prodrug (S1) | Ac-AKRHHGYKRKFHGGVKKK/DVVDGGDDDDDD | 3434.6778 | 0 |
| *P. gingivalis*<br>prodrug (G1) | AKRHHGYKRKFHGGAGPR/IDGGDDDDDD | 3090.4474 | 0 |

The second part (**Scheme 1** green) blocks the antimicrobial activity before cleavage via a poly-aspartic acid sequence (DDDDDD). These negative charges balance the positive charge of P-113 creating a zwitterionic peptide that is non-toxic until protease release separates the therapy from the quencher.

The third part (**Scheme 1** blue) is the tunable cleavage sequence and makes the prodrugs selective to *C. albicans* or *P. gingivalis*. *C. albicans* releases a portfolio of proteases, but we chose SAP9 because it is representative of the *Candida* genus.[33] SAP9 cleaves primarily after basic amino acids such as lysine[15]—particularly a triple lysine residue. We thus chose the substrate <u>VKKK/DVVD</u>.[30] For *P. gingivalis*, we used arginine-gingipain B (RgpB)—an endopeptidase that can cleave peptides on the C-terminal of arginine.[18] This protease cleaves between arginine and isoleucine (Arg/Ile), and <u>AGPR/ID</u> is a substrate with a high affinity for RgpB.[18] Hence, the specific substrates for either SAP9 or RgpB were incorporated into the peptide design. Di-glycine spacers (GG) were placed between the three different functional parts to provide flexibility. We defined the *C. albicans* prodrug as S1 and the *P. gingivalis* prodrug as G1; see Table 1 for full sequences.

### Probe activation, kinetics, and cytotoxicity

P-113, S1, and G1 were synthesized through standard solid-phase Fmoc synthesis, purified using high-performance liquid chromatography (HPLC), and characterized using matrix-assisted laser desorption ionization–time-of-flight (MALDI-TOF) mass spectrometry (see Methods). The cleavage of S1 by SAP9 (300 nM) and G1 by RgpB (200 nM) was first investigated *in vitro* using recombinant proteases via a 3-hour incubation at 37 °C in the appropriate activity buffer as recommended by previous work (9.5 mM MES, 2.7 mM KCl, 140 mM NaCl, pH 5.5 for SAP9 and 0.2 M Tris HCl pH 7.6, 5 mM CaCl_2_, 100 mM NaCl and 40 mM TCEP for RgpB).[30, 34] MALDI-TOF showed obvious mass peaks corresponding to the cleaved fragments (**Figure 1B-C**). As a control, P-113 (**Supplemental Figure S2**) was incubated with the proteases, but mass spectrometry showed no degradation.

To evaluate the cleavage efficiency of SAP9, a fluorescent analogue of S1 was synthesized (**Figure 1D, Supplemental Figure S3**) using Cy5.5 and Cy3, thus producing strong FRET quenching. Once cleaved by SAP9, the dyes were separated for increased fluorescent signal. This probe was incubated with 200 nM of the recombinant enzyme at 37 °C and fluorescence was monitored longitudinally to calculate the k_cat_/K_M_ value as ∼55,000 M^−1^ s^−1^ (**Supplemental Figure S4**).[43] The cleavage efficiency of RgpB is known from previous work to be 12,000 M^−1^ s^−1^.[44] [45]. The higher k_cat_/K_M_ ratio for SAP9 suggests that SAP9 works better with less substrate, thus enabling it to cleave the peptide more readily (**Figure 1E**).

To verify the prodrugs’ antimicrobial activity and proteolytic selectivity, we first grew *C. albicans* and *P. gingivalis* from a lyophilized stock. The presence of secreted aspartyl protease was confirmed via a commercially available ELISA kit as shown in **Supplemental Figure S5**. RgpB secretion was confirmed similarly as described in previous work.[18] Next, a Kirby-Bauer disk diffusion assay was conducted to evaluate initial sensitivity of the microorganisms to the prodrugs. For *C. albicans*, a clear diameter was formed by addition of S1 to the agar showing inhibited growth of the fungus to approximately 70% of the positive control treatment (**Figure 2A-B)**. Alternatively, the negative control showed colony growth around and under the disk, thus demonstrating a lack of antimicrobial activity as presented quantitatively (**Figure 2C**). Similarly, G1 created a 10-mm diameter on the *P. gingivalis* plate, nearly half that of the antibiotic treatment (**Figure 2D-F**). In comparison, the negative control showed no measurable diffusion of *P. gingivalis* colonies, with rich colony formation near the treatment area. These disk diffusion tests confirm that the prodrug could inhibit the growth of the respective microorganism.

**Figure 2.**
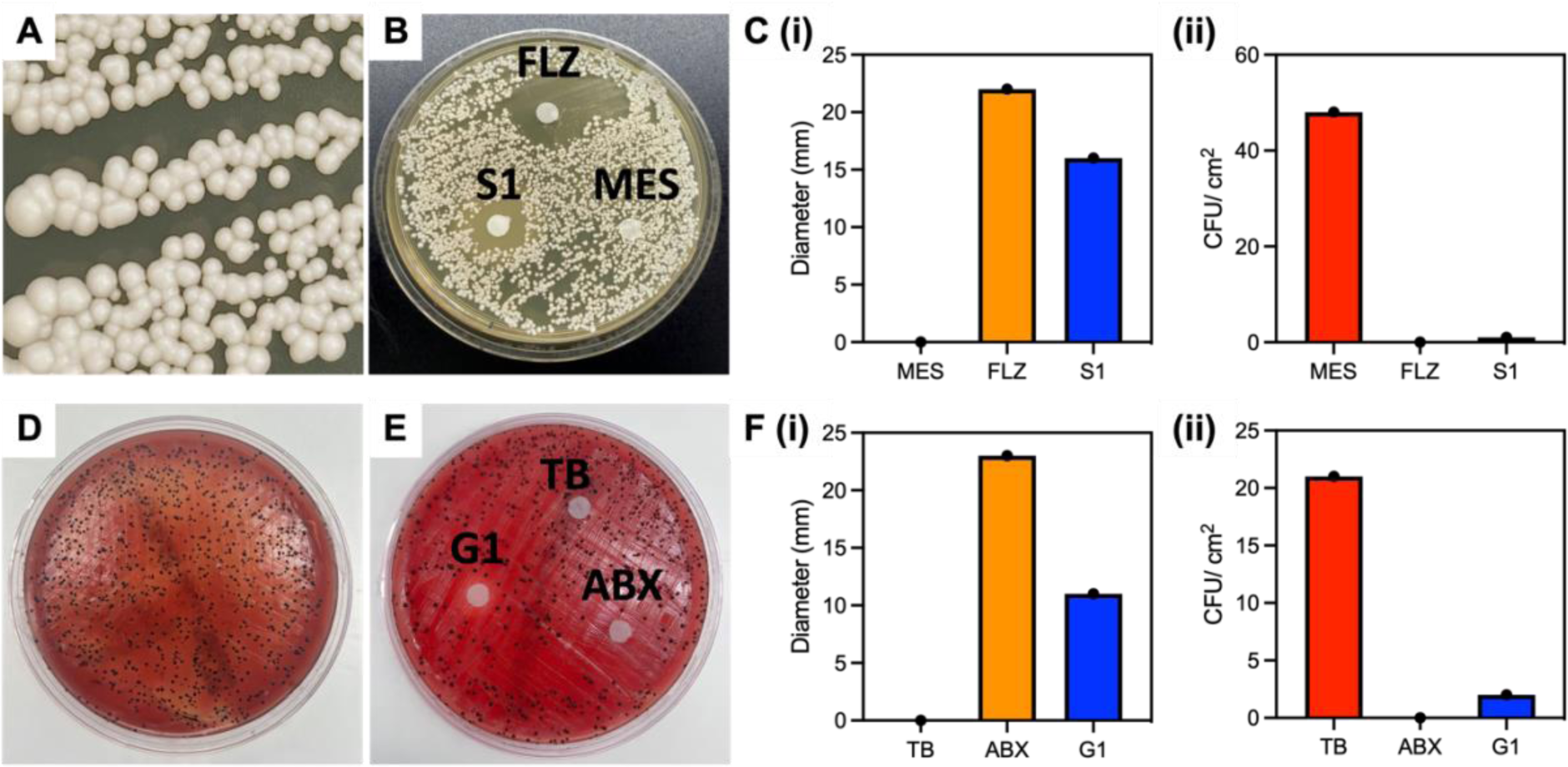
Microorganism growth and disk diffusion test. **(A)** “String of pearls” characteristic of *C. albicans* formation. **(B)** *C. albicans* disk diffusion test after five-day incubation at 30 °C with MES buffer (negative control), 1 mM S1 prodrug, and 1 mM Fluconazole (positive control) disks showing a clear growth diffusion with the experimental and positive control. Conversely, the negative control shows no inhibition of growth as represented by colonies forming around and beneath the treated filter. **(C) (i)** Quantitative diameter measurement for three disks in **Figure 2B** and **(ii)** colony forming units surrounding the disk per 1 cm^2^. **(D)** *P. gingivalis* colonies with characteristic black pigmentation plated and grown anaerobically on blood agar. **(E)** *P. gingivalis* disk diffusion test with sterile Tris buffer (negative control), 1 mM G1 prodrug, and 1 mM penicillin-streptomycin antibiotic (positive control) showing diffusivity diameter among experimental and positive control disks compared to the buffer with colonies growing beneath and around the disk. **(F) (i)** Quantitative diameter measurement for three disks and **(ii)** colony forming units surrounding the disk per 1 cm^2^.

Viability assays were used next to quantitate cleavage-dependent toxicity against *C. albicans* and *P. gingivalis* as well as two control cell lines: *Fusobacterium nucleatum* and human embryonic kidney cells HEK293T (**Table 2**). *F. nucleatum* is an interesting negative control because it is also present in the gingival sulcus but is commensal and not known to produce either secreted aspartyl proteases or gingipains.[18] S1 and G1 were expected to show low toxicity due to the absence of specific cleavage. Mammalian cells were unlikely to be targeted when exposed to the prodrugs due to differences in their membrane structure and composition relative to fungi and bacteria. We selected HEK293T as model mammalian cells.[46]

**Table 2.** Key microorganisms investigated.

| Microorganism | Target Protease | Features |
| --- | --- | --- |
| *C. albicans* | SAP9 | S1 peptide target |
| *P. gingivalis* | RgpB | G1 peptide target |
| *F. nucleatum* | - | Cleavage negative control |
| HEK 293T | - | Toxicity negative control |

**Figure 3A (i)** shows the cytotoxicity of 10 µM P-113, PBS, S1, and G1 against the different cell lines outlined in **Table 2**. The incubation time was optimized to 3 hours to allow for successful cleavage and therapeutic release. Excess time (> 6 hours) led to prodrug degradation and eventually (> 9 hours) cellular regrowth as shown by a decrease in percent toxicity. Similarly, insufficient time (< 3 hours) resulted in a decreased toxic effect, suggesting a lower product turnover rate of the prodrugs (**Supplemental Figure S9**). These findings are consistent for both S1 and G1. As expected, P-113 was only toxic to the oral microorganism cell lines at an average 50% toxicity rate and demonstrated slight toxicity towards HEK293T at 23%, never exceeding 50% even at 100 µM treatment.

**Figure 3.**
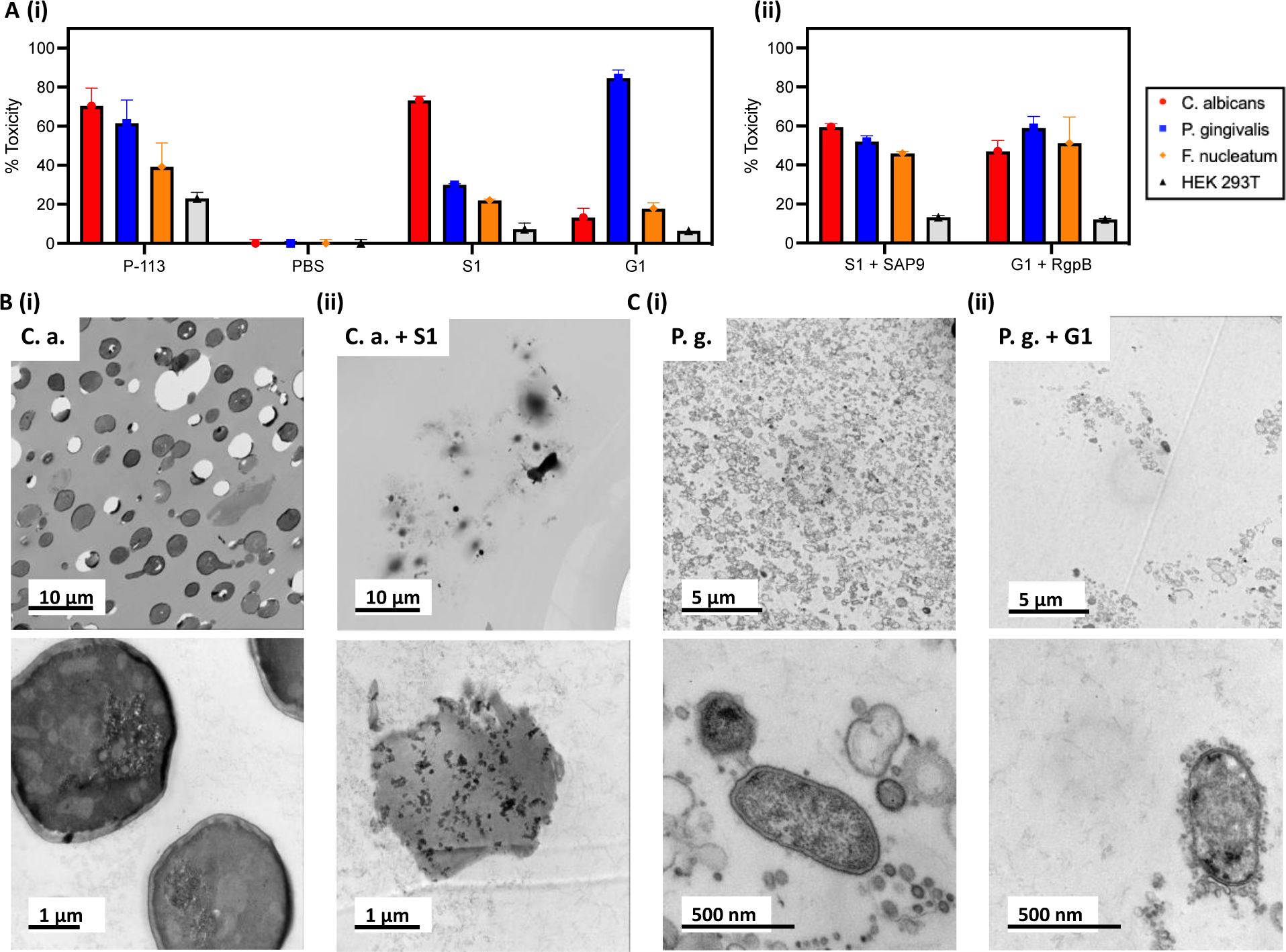
Influence of prodrugs on oral microorganisms and healthy mammalian cells. **(A) (i)** XTT viability assay at 10 µM treatment of P-113 (positive control), PBS (negative control), S1 (*C. albicans* prodrug), and G1 (*P. gingivalis* prodrug) in four cell lines: *C. albicans*, *P. gingivalis*, *F. nucleatum*, and HEK 293T. Percent toxicity shows that P-113 has an increased effect on oral microorganisms. Conversely, S1 has a negligible effect on all cell lines apart from that which secrete SAP9 (*C. albicans*). Similarly, G1 only exerts a toxic effect greater than 50% against *P. gingivalis*. **(ii)** Viability assay for S1 and G1 cleaved by 300 nM or 200 nM, respectively, of recombinant proteases *in vitro* prior to cell treatment. Here, a return in toxicity similar to that exerted by P-113 is prevalent in all three oral microorganisms. Still, a decreased toxicity is exerted upon HEK293T. **(B)** and **(C)**Transmission electron microscopy (TEM) of 70 nm thin sections of plastic embedded samples to visualize cellular organization. **(B)** Scale bar presents 10 µm (top) and 1 µm (bottom). **(i)** *C. albicans* (**C.a**.) cells alone, showing dense cellular population at low magnification and rigid, intact cell membrane at high magnification. **(ii)** *C. albicans* cells with 40 µM S1 treatment after incubation for 3 hours at 37 °C (**C. a. + S1**) demonstrating consequential decreased cell density and cell rupturing. **(C)** Scale bar presents 5 µm (top) and 500 nm (bottom). **(i)** *P. gingivalis* (**P.g**.) cells alone showing full growth and clear in-tact outer membrane. **(ii)** *P. gingivalis cells* with 40 µM G1 treatment after incubation for 3 hours at 37 °C (**P.g. + G1**) demonstrating overall cell density decrease at 5 µm (top) and cell-membrane rupture and leakage at 500 nm scale (bottom). Different magnifications are shown in Supplemental Information.

The minimum inhibitory concentration (MIC) was an average of 5.92 µg/mL for P-113 against the three oral microorganisms and nearly 100 times that for HEK293T.[47] This confirms the charge dependence of the P-113 cytotoxic mechanism. *C. albicans* showed increased susceptibility compared to the bacteria. S1 and G1 were toxic to *C. albicans* (70%) and *P. gingivalis* (52%) at 10 µM, consistent with a MIC of 3.4 and 3.1 µg/mL, respectively. Alternatively, the toxicity of S1 and G1 against *F. nucleatum* never exceeded 50% and showed slight inhibition (15%) at 3 µg/ mL. The prodrugs, S1 and G1, also showed a decreased toxicity by nearly half to HEK293T, compared to P-113, suggesting that the intact zwitterionic structure reduced toxicity to mammalian cells. Importantly, when S1 was pre-cleaved *in vitro* by 300 nM recombinant SAP9 and G1 by 200 nM recombinant RgpB, the resulting compounds expressed toxicity towards the three oral microorganisms (**Figure 3A(ii)**). This, in turn, confirms that once cleaved, the prodrugs S1 and G1 release the fragment responsible for cell death. Importantly, the toxicity of the prodrug has nearly molar equivalency to the parent P-113 AMP in the target microorganism suggesting a complete return after cleavage. Full titration curves are in **Supplemental Figure S10-S14**.

The target microorganisms were also imaged using transmission electron microscopy (TEM) to visualize the cell morphology. **Figure 3B (i)** presents *C. albicans* cells after the standard growth protocol. The membrane structure is rigid and intact in contrast to **Figure 3B (ii)** showing decreased cell density and clear destruction of the membranes. This obvious change in morphology is seen at other magnifications (**Supplementary Figure S15-S16**). Similarly, **Figure 3C (i)** shows untreated *P. gingivalis* with characteristic gram-negative outer membrane versus treated bacteria in **Figure 3C (ii)** with decreased cell wall density and cellular leakage characteristic of apoptosis. Confocal microscopy was used to image HEK293T to ensure that the membrane remained intact after treatment by S1 and G1 (**Supplementary Figure S7**).

### Colorimetric Detection of Pathogens

Beyond therapy, this same system could offer diagnostic utility. Detection of high SAP9 and RgpB levels is important because it is a direct marker of pathogenesis with few convenient detection methods—especially chairside methods for dental professionals.[48] Colorimetric detection via plasmonic particles offers a convenient, point-of-care advantage over conventional or symptomatic diagnoses.[35, 49, 50] Herein, we investigated the controlled assembly of gold (AuNPs) and silver (AgNPs) nanoparticles.

For gold, citrate-capped AuNPs (15 nm diameter) were synthesized and had a negatively charged coating.[51] AgNPs (20 nm diameter) were coated with bis(p-sulfonatophenyl)phenylphosphine (BSPP) via a seed-mediated growth procedure.[52] Here, a BSPP coating was primarily chosen for its negative charge. We previously demonstrated that the BSPP coating on silver, unlike citrate, can desorb from the AgNPs during positively charged peptide-mediated assembly, thus releasing Ag+ into the solution and permitting spontaneous assembly of the particles. Both AuNP-citrate and AgNP-BSPP possess a distinct wavelength of light absorption as well as an associated extinction coefficient. The assembly for the nanoparticle systems is thus promoted via electrostatic interaction between the negatively charged particle surface and the positively charged fragment released after proteolytic cleavage (**Scheme 2**).

**Scheme 2.**
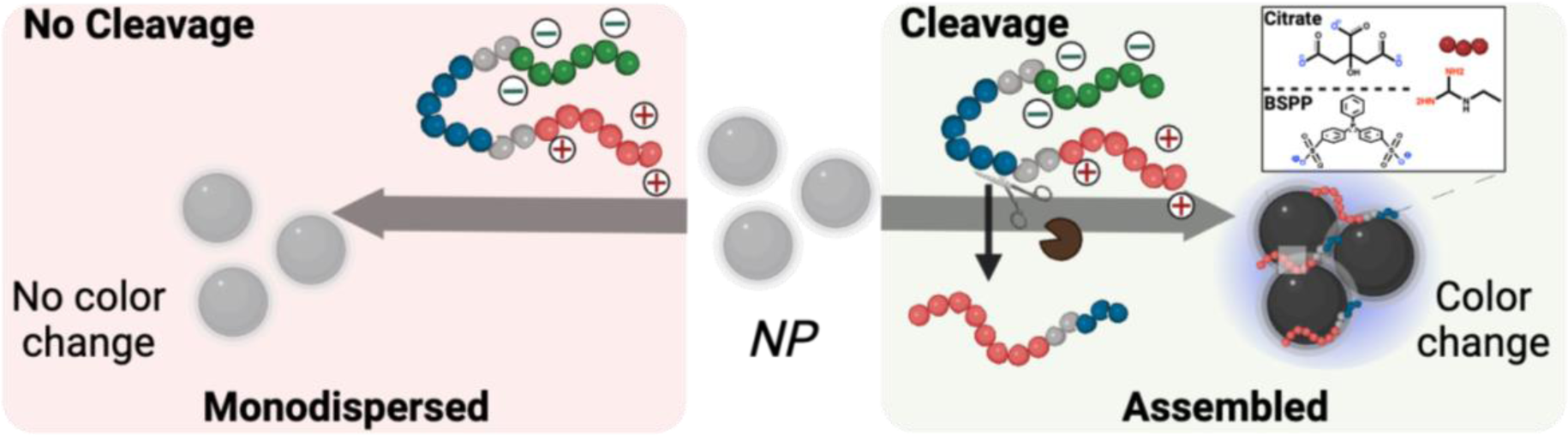
Descriptive schematic of enzymatic detection with plasmonic nanoparticles. Color evolution of monodispersed coated-nanoparticles in the presence of intact zwitterionic prodrug (left) vs. the cleavage-induced assembly of the nanoparticles caused by pathogenic enzymatic activity. Citrate-coated gold nanoparticles, (right inset, top) or BSPP-coated silver nanoparticles (right inset, bottom) were used as independent colorimetric systems for comparison. In turn, the cleavage of the prodrug results in a color change from ruby red to deep blue for AuNPs and yellow to bright blue for AgNPs.

First, the AuNP-citrate were mixed with S1 and G1, independently. The zwitterionic design is interesting for controlled assembly of plasmonic nanoparticles because the charge-shielding site effectively quenches the positively charged assembling site prior to proteolytic activity within the cleavage substrate. The colloidal state of the AuNPs-citrate suspension was defined as the ratio of the absorbance: A_700 nm_/A_520 nm_.[36, 37] High and low ratios correspond to assembled or monodispersed NPs, respectively. The maximum ratio was obtained for concentrations of 25 µM and 5 µM of S1 and G1, respectively. However, when S1 and G1 were cleaved by recombinant proteases, the concentration needed to induce the assembly dropped to 1.5 and 0.4 µM for S1 and G1, respectively, showing that the colloidal state is responsive to the biomarker activity (**Figure 4A**). Visually, the AuNP-citrate suspensions turned from bright red to blue almost instantly after addition in the presence of the proteases (**Figure 4B**). AgNP-BSPP were used similarly wherein the colloidal state of the suspension was defined as the ratio of the absorbance : A_650 nm_/A_400 nm_.[52] Here, the maximum ratio was obtained at 25 µM for both S1 and G1. However, when S1 and G1 were cleaved with the corresponding protease, the concentration needed to assemble the silver nanoparticles dropped to 6 µM and 0.4 µM, again showing that the colloidal state is responsive to the protease activity (**Figures 4C**). Similarly, visual detection was again clear for silver in a transition from yellow to purple (**Figure 4D**). This was repeated in saliva (**Supplemental Figure S20**) to demonstrate the applicability of this detection system wherein 10% saliva returned a similar trend. The assembly was further verified through TEM (**Figure 4E-F**) demonstrating the transition from monodispersed to peptide-assembled as a consequence of the cleavage of the prodrugs by the proteases.

**Figure 4.**
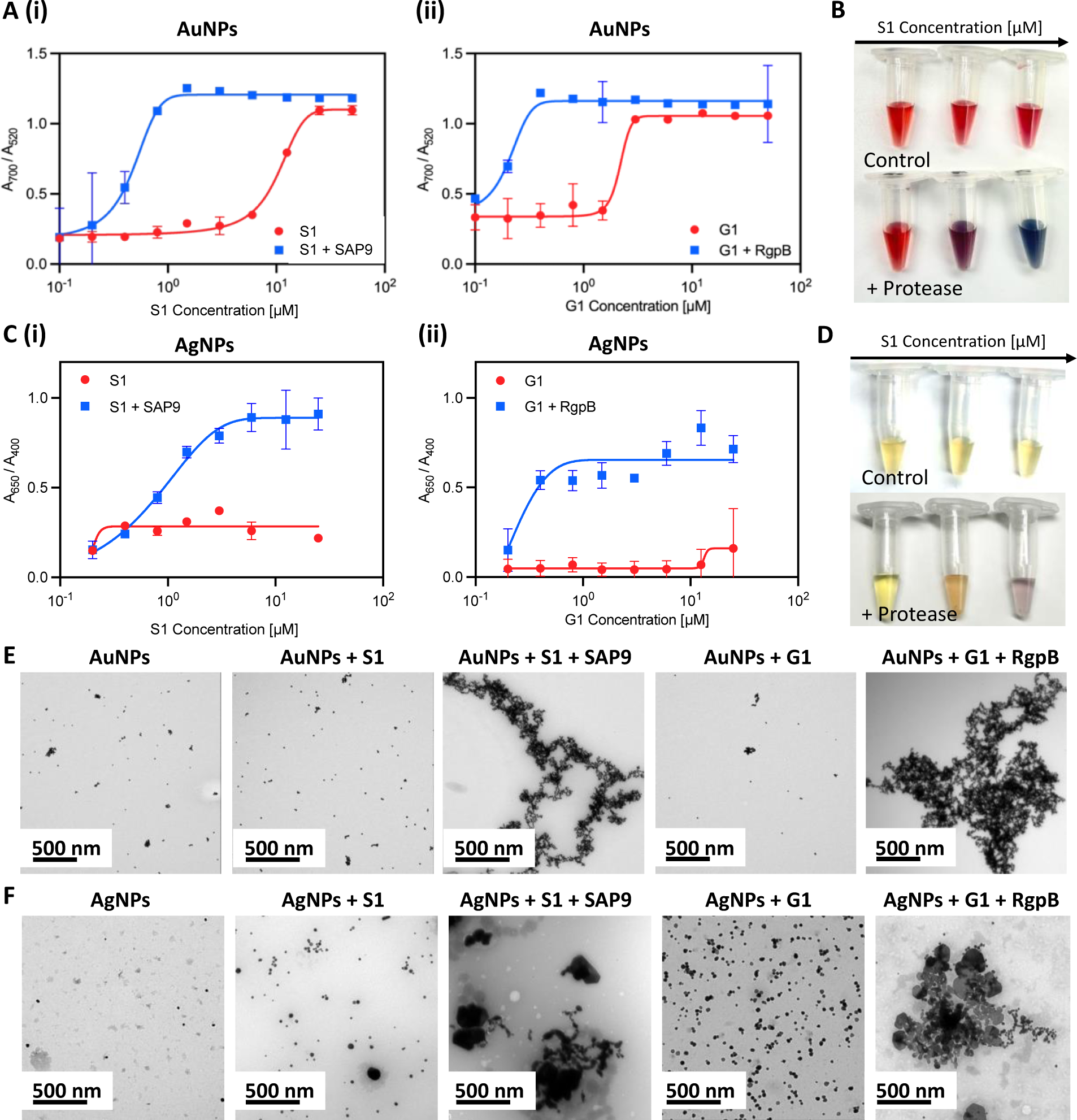
Protease-induced plasmonic nanoparticle assembly. **(A)** Optical absorption of AuNPs when incubated with increased concentrations of **(i)** S1 and **(ii)** G1 parent (red) and fragmented peptides (blue). **(B)** The color evolution of AuNPs in the presence of intact S1 (control) and cleaved S1 by recombinant SAP9 in increasing concentration from 0.01 to 10 µM wherein high concentration of cleaved peptide results in a transition from ruby red to deep blue, indicative of increased particle assembly caused by isolated positive fragments post-cleavage. **(C)** Optical absorption of AgNPs when incubated with **(i)** S1 and **(ii)** G1 parent and fragmented peptides. **(D)** The color evolutions of AgNPs in the presence of intact S1 (control) and cleaved S1 by recombinant SAP9 in increasing concentration from 0.01 to 10 µM. Color change from yellow to purple represents increased particle assembly. **(E)** Corresponding TEM images confirming monodispersion or assembly of AuNPs and **(F)** AgNPs when treated with PBS (**NPs**), intact S1 (**NPs + S1**), cleaved S1 (**NPs + S1 + SAP9**), intact G1 (**NPs + G1**) and, cleaved G1 (**NPs + G1 + RgpB**) (left to right).

Next, the peptides were investigated together in solution. The goal was to optimize the system to address both *C. albicans* and *P. gingivalis* for a dual sensing mechanism. The peptide-duo was subjected to four conditions: no enzyme, SAP9, RgpB, and an equimolar combination of SAP9 and RgpB. Here, the two peptides were incubated in solution (**Figure 5A**) and the cleavage of the mixed peptides was reevaluated using MALDI-TOF. The cleavage confirmation was presented as previously, based on the emerging peaks indicative of the fragment mass. A clear peak for G1 remained present during incubation with SAP9 (**Figure 5B**). Likewise, S1 remained intact when the duo-peptide solution was incubated with RgpB, thus demonstrating the affinity of the proteases to their cleavage substrate (**Figure 5C**). Only during the coincubation with both proteases were both prodrugs cleaved as indicated by the fragment peaks on the spectra (**Figure 5D**).

**Figure 5.**
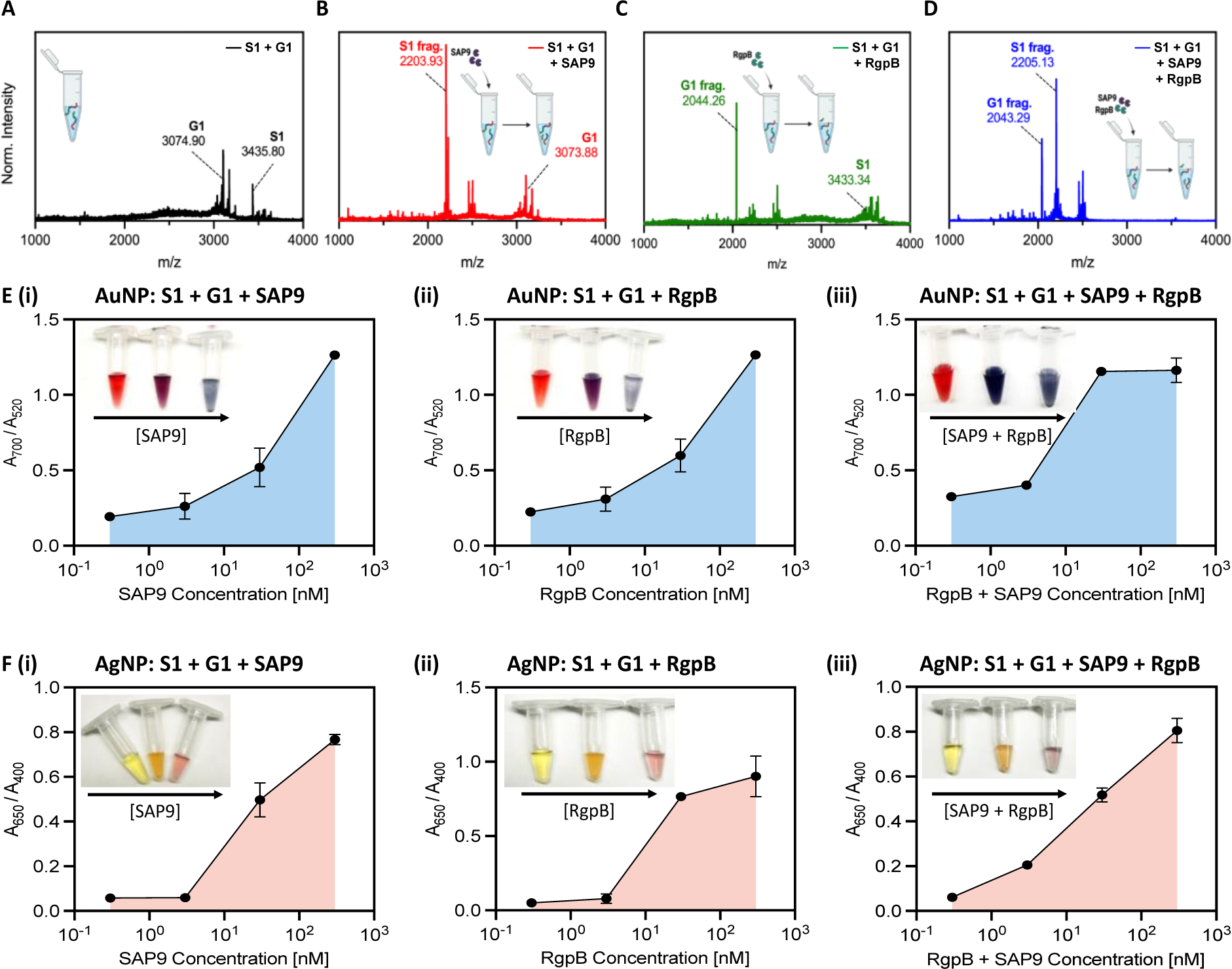
Combined cleavage and limit of detection. MALDI-TOF of (A) joint incubation of intact S1 and G1 peptides showing successful synthesis and co-incubation. (B) S1 and G1 incubated with SAP9 (300 nM, 3 hours, 37 °C), showing S1 converting to S1 fragment (2203 Da), and intact G1, with no peak corresponding to a G1 fragment. (C) S1 and G1 incubated with RgpB (300nM, 3 hours, 37 °C), showing S1 remaining intact, and G1 cleaving as confirmed by the G1 fragment peak (2044 Da). (D) S1 and G1 incubated with 300 nM SAP9 and RgpB for 3 hours at 37 °C, showing both prodrugs cleaving as presented by only fragment peaks. (E) Limit of detection (LOD) for AuNP-citrate and (F) AgNP-BSPP detection system of co-incubated S1 and G1 using an increasing concentration of (i) SAP9, (ii) RgpB, and (iii) both, SAP9 and RgpB. LOD study shows a decrease in the limit for both proteases and a detectable limit within the regime of physiologically relevant protease secretion concentrations. Inset presents color evolution at 0 nM, 3 nM, and 300 nM (left to right).

The peptides were next subjected to a colorimetric assay with AuNPs at varying concentrations of the enzymes. A limit of detection (LOD) of 10 nM (**Figure E, i**) was obtained for SAP9, which agrees well with previously reported protease detection among similar systems.[18, 35, 36] The color change was also clear as an increased enzymatic concentration returned a progression from red to purple and finally, blue. RgpB yielded a similar LOD with the gold system at approximately 10 nM (**Figure E, ii**). Conversely, when both enzymes were incubated in the duo-peptide solution, the LOD was nearly halved as suggested by the more intense color change and enthusiastic assembly of the particles. Here, 3 nM protease treatment resulted in a deep blue color versus the previous purple color (**Figure E, iii**).

The silver system proved more sensitive, with a LOD of approximately 3 nM for SAP9 and RgpB (**Figure F, i-ii**). The color change was obvious by eye from yellow to purple. The system was more responsive when both proteases were present with a LOD of 0.3 nM. The color change followed in a more dramatic change from yellow to purple-blue (**Figure F, iii**). This data further suggests that due to their ability to produce a stronger plasmon resonance than gold, silver is able to absorb light more readily making it more sensitive to peptide assembly. Moreover, the observed tenfold increase in sensitivity of the silver system is consistent with previous work.[34]

Further, this limit was assessed in saliva (**Supplemental Figure S21**). Here, the LOD increased by a factor of 10 for both enzymes. It is likely that saliva interferes with nanoparticle sensors by suppressing color change signals due to nonspecific protein absorption.[53] As such, this increase is expected. Another possible explanation for this is the negatively charged proteins forming a protein corona around the nanoparticles thus impeding the aggregation.[54] It is, however, worth noting that several factors influence the LOD, including the specificity of the detection method, the affinity of the probe for the target protease, the signal amplification strategies employed, and the background noise in the system. The variation between enzymatic LOD for SAP9 and RgpB can be credited to the higher kcat/KM value of SAP9. As such, given the kinetic information of the design and the low MIC yielded, the LOD is satisfactory and well in line with the physiological concentration of SAP9/RgpB.[44, 55–57]

## CONCLUSION

We report a therapeutic and diagnostic scaffold for *C. albicans* and *P. gingivalis* that harnesses the value of antimicrobial peptides for oral health. Although P-113 has been pursued for years due to its antimicrobial properties, the activatable form is a completely novel approach to the development of antimicrobial therapies. By offering specific release, the prodrug-peptides demonstrated high toxicity (MIC = 3 µg/ mL) towards the microorganisms expressing the trigger protease. Alternatively, when subjected to conditions absent of the target, the prodrugs delivered relatively no toxicity and never exceeded 50% cell death even at concentrations as high as 300 µg/ mL. Finally, the charge-switchable design was exploited for *in vitro* colorimetric detection using plasmonic nanoparticles, a feature of P-113 that had yet to be explored. Validation of this system demonstrated rapid detection of the relevant toxicity markers of *C. albicans* and *P. gingivalis* at nanomolar enzymatic concentrations. Importantly, this combination theragnostic approach is promising for developing controlled preventative treatment and enhanced diagnoses in the oral microbiome. It is worth noting that increasing the affinity of the proteases to the cleavage substrate can improve the design by increasing the effectiveness of the therapy and limit of detection. However, we imagine that this prodrug strategy can be extended to any enzyme after identifying the appropriate peptide substrate making a sustainable approach to theranostic devices.

## EXPERIMENTAL SECTION

### Enzyme digestion

SAP9 and RgpB cleavage were both performed with 455 µM of peptide. Typically, 50 µL of peptide (1 mM) were mixed with 50 µL of buffer (9.5 mM MES, 2.7 mM KCl, 140 mM NaCl, pH 5.5 for SAP9 and 0.2 M Tris HCl pH 7.6, 5 mM CaCl_2_, 100 mM NaCl and 40 mM TCEP for RgpB). Then, 10 µL of the enzyme at the appropriate concentration was added. An optimized final concentration of 300 nM for SAP9 and 200 nM for RgpB were used. The solution was incubated at 37°C for 3 hours for RgpB and SAP9 for optimal cleavage.

### Fungal culture

*In situ* culturing *C. albicans* (ATCC 90028) from lyophilized stock (KWIK-STIK, VWR) was conducted on Sabouraud dextrose agar plates (Sigma-Aldrich) 2 to 5 days in advance of the antifungal assay.[58] The plates were then incubated at 30°C and pearl-like colonies were observed, indicative of *C. albicans*. Single colonies were then extracted with sterilized inoculation loops and inoculated in 3 mL yeast extract peptone dextrose (YEPD) medium (2% Bacto^TM^peptone, 2% Dextrose, 1% Yeast extract in H_2_O) using Biological Safety Level 2 (BSL-2) practices and procedures. Note, the broth was autoclaved before use and stored at room temperature. The inoculates were then grown overnight with gentle shaking at 30 °C. The culture was finally diluted 1:100 and the OD600 was measured wherein OD600 = 0.5 a.u. correlates to approximately 1 × 10^6^ cells/ mL in 100 µL RPMI-1640 medium with L-glutamine and 3-(N-morpholino)propane sulfonic acid (MOPS) without sodium bicarbonate, pH 7.0. Next, the 96-well plates were sealed with a sealing membrane to reduce evaporation and prevent cross contamination before being incubated at 37°C to allow biofilms to form overnight. The media was then carefully discarded, and the biofilms were washed from free cells with 200 µL sterile PBS.

### Bacteria culture

*P. gingivalis* and *F. nucleatum* colonies were grown on brucella blood agar plates (Sigma) from lyophilized stocks of ATCC 33277 and ATCC 25586, respectively (KWIK-STIK, VWR).[18] Anaerobic atmospheres were generated and maintained using anaerobic bags with oxygen indicators (GasPak EZ Pouch System, BD). Briefly, the plates were streaked with bacteria-loaded swabs and immediately placed in the gas-generating sachets. For *P. gingivalis*, light-colored colonies became visible 4-5 days of incubation at 37°C; these colonies turned black over the following four days due to heme accumulation/metabolism (signature of *P. gingivalis*). *F. nucleatum* colonies were faster growing (2-4 days) and maintained a light color. Single colonies were transferred to enriched tryptic soy broth (eTSB) which was prepared at 500-mL scale by dissolving 15 g tryptic soy broth and 2.5 g yeast extract in 450 mL DI water followed by addition of 2.5 mL hemin (1 mg/mL). The pH was adjusted to 7.4 using 5 M NaOH and water was added to reach 500 mL. The broth was then autoclaved followed by aseptic addition of 5% (w/v) L-cysteine, 10 mL 1% (w/v) dithiothreitol, and 2 mL menadione (0.5 mg/mL).[18]

### Viability assay

The percent toxicity was calculated using an XTT (2,3-bis-(2-methoxy-4-nitro-5-sulfophenyl)-2H-tetrazolium-5-carboxanilide) colorimetric assay (Biotium) wherein viable cells form an orange formazan product upon reduction. Here, the oral microorganism cells were grown under appropriate conditions until the desired growth phase of 1 × 10^6^ cells/ mL in a 96-well plate. Control wells with medium only served as the blank control and wells with cells treated with fluconazole or penicillin-streptomycin agent served as the positive control. Finally, an increasing concentration of the various peptides were used to treat the cells in sextuplet replicate measurements. The plates were incubated at 37 °C for 3 hours allowing the antifungal agents to exert their effects. XTT solution was then added to each well as recommended by the provider. The plates were then incubated in the dark at the appropriate temperature for 3 hours. The absorbance of the formazan dye was measured at a wavelength of 490 nm using a microplate reader. Here, the absorbance is directly proportional to the number of viable cells present in each well. Thus, higher absorbance indicates higher cell viability. The readings were then normalized by subtracting the average absorbance of the blank control wells. Finally, the percentage of cell viability for each treatment group was calculated by comparing the absorbance of treated wells to the control wells. The MIC was also determined after subtraction of the blank control consisting of growth medium only. Here, readings below 0.01 optical density units (i.e., no visible growth) indicated growth inhibition. Independent MIC determinations with the same isolate and compound were usually identical or within a twofold range. The average was collected along sextuplet measurements collected on two independent plates.

### AuNP synthesis

Citrate-stabilized AuNPs were synthesized at 15 nm using the Turkevich method.[51] Briefly, 45 mg of HAuCl_4_·3H_2_O was dissolved in 300 mL of MQ water while subjected to generous stirring (600 rpm) and boiling conditions at 120 °C via oil bath. Next, 150 mg of sodium citrate (dissolved in 5 mL of MQ water) was rapidly injected, and the reaction was left under boiling conditions for 20 minutes. The color of solution changed from purple to gray and finally dark red. The resulting product was cooled and stored at room temperature for the future use. The optical density of the final product was 1.45 (concentration ∼ 3.6 nM, ɛ_520_ = 4.0 × 108 M^−1^cm^−1^). Notably, the flask used was cleaned thoroughly with Agua regia and distilled water (three times) prior to synthesis.

### AuNP detection

100 µL of 15 nm AuNPs-citrate (OD = 1) were mixed with varying concentration of the peptides in buffer, allowing the color of the suspension to turn from bright ruby-red to purple, and ultimately, deep blue.

### AgNP synthesis

BSPP-coated AgNPs (AgNPs-BSPP) were synthesized using a two-step procedure.[52] First, silver seeds were produced by the reduction of silver nitrate (0.1 mM, 6 mL) with sodium borohydride (0.1 M, 60 μL) in a glass vial under magnetic stirring for 16 h. The seeds were then grown into nanoparticles by adding sodium ascorbate (15 mM, 400 μL), BSPP (5 mM, 200 μL), dropwise silver nitrate (1 mM, 4 mL), and then BSPP again (5 mM, 400 μL), and the resulting solution was stirred for 48 h at room temperature.

### AgNP detection

100 µL of 20 nm AgNPs-BSPP (OD = 1) were mixed with increasing concentrations of the peptides in aqueous solution, allowing the color of the suspension to turn from bright yellow to orange, red and ultimately, bright blue.

## ASSOCIATED CONTENT

### Supporting Information

Materials, experimental methods, instrumentation, synthesis characterization, microscopy magnification variation, and expanded toxicity assay titrations.

## ACKNOWLEDGMENT

This work was sponsored by the UC San Diego Materials Research Science and Engineering Center (UCSD MRSEC), supported by the National Science Foundation (Grant DMR-2011924). The authors thank the University of California, San Diego - Cellular and Molecular Medicine Electron Microscopy Core (UCSD-CMM-EM Core, RRID: SCR_022039) for equipment access and technical assistance. The UCSD-CMM-EM Core is partly supported by the National Institutes of Health Award number S10 OD023527. We also acknowledge the UCSD School of Medicine Microscopy Core (Grant P30 NS047101) and NIH funding via R21 DE029917 and R01 DE031307.

