## Supplemental Data for "Activatable prodrug for controlled release of an antimicrobial peptide via the proteases overexpressed in *Candida albicans* and *Porphyromonas gingivalis*"

Table of Contents


I. Materials ____ 3

II. Experimental Procedures ____ 3

III. Supplementary Data ____ 6

Figure S1. Structure of P-113 and SAP9 and RgpB- cleavable prodrugs, S1 and G1, respectively (top to bottom). ____ 6

Figure S2. P-113 characterization. ____ 7

Figure S3. SAP9 cleavable substrate FRET probe characterization. ____ 8

Figure S4. SAP9 cleavable substrate FRET probe fluorescence. ____ 9

Figure S5. ELISA detection of SAP in *C. albicans* culture. ____ 10

Figure S6. *C. albicans* growth transmission microscopy. ____ 11

Figure S7. Confocal microscopy of mammalian cells. ____ 12

Figure S8. Transmission, NucBlue, and propidium iodide microscopy for *C. albicans*. ____ 12

Figure S9. Time dependent toxicity assessment for S1 and G1. ____ 13

Figure S10. Toxicity assessment for varying concentrations of P-113, positive control, treatment along four cell lines. ____ 14

Figure S11. Toxicity assessment for varying concentrations of S1, C. albicans prodrug, treatment, along four cell lines. ____ 15

Figure S12. Toxicity assessment for varying concentrations of G1, P. gingivalis prodrug, treatment, along four cell lines. ____ 16

Figure S13. Toxicity assessment for varying concentrations of S1 + SAP9, C. albicans prodrug pre-cleaved by 200 nM recombinant protease in vitro, treatment, along four cell lines. ____ 17

Figure S14. Toxicity assessment for varying concentrations of G1 + RgpB, P. gingivalis prodrug pre-cleaved by 200 nM recombinant protease in vitro, treatment, along four cell lines. ____ 18

Figure S15. C. albicans TEM images of cellular morphology at increased magnification. ____ 19

Figure S16. P. gingivalis TEM images at increased magnification. ____ 20

Figure S17. Heat map for varying protease concentration in LOD analysis. ____ 21

Figure S18. Colorimetric detection titration curves for combined S1 and G1. ____ 21

Figure S19. Gram-positive bacteria toxicity assay of P-113 against MRSA USA300. ____ 22

Figure S20. Protease-induced plasmonic nanoparticle assembly in saliva. ____ 23

Figure S21. Limit of detection of plasmonic nanoparticle assembly in saliva. ____ 24

IV. Supplementary References ____ 25

V. Author Contributions ____ 25

### I. Materials

Fmoc-protected L-amino acids, hexafluorophosphate benzotriazole tetramethyl uranium (HBTU), and Fmoc-Rink amide MBHA resin (0.67 mmol/g, 100-150 mesh) were purchased from AAPPTec, LLC (Louisville, KY). Organic solvents including N,N-dimethylformamide (DMF, sequencing grade), acetonitrile (ACN, HPLC grade), ethyl ether (certified ACS), methylene chloride (DCM, certified ACS), and dimethyl sulfoxide (DMSO, certified ACS) were from Fisher Scientific International, Inc. (Hampton, NH). Ultrapure water (18 MΩ.cm) was obtained from a Milli-Q Academic water purification system (Millipore Corp., Billerica, MA).

Cy5.5-NHS and Cy3-NHS were purchased from Lumiprobe, Inc. (Maryland, USA). Dimethyl sulfoxide, leupeptin, phosphate-buffered saline, tris base/HCl, dithiothreitol, triethylamine, L-cysteine, menadione, yeast extract, tryptic soy broth, dextrose, and hemin were purchased from Sigma-Aldrich (Missouri, USA). Acetonitrile, trifluoroacetic acid, RPMI-1640 medium, 3-(N-morpholino)propane sulfonic acid, Bacto™peptone, and sodium hydroxide were purchased from Fisher Scientific (Massachusetts, USA). Recombinant SAP9 and SAP ELISA Kit were purchased from MyBioSource (San Diego, CA). Recombinant RgpB was a kind gift from Professor Anthony O'Donoghue (UC San Diego Skaggs School of Pharmacy and Pharmaceutical Sciences, San Diego, CA).

*C. albicans* (ATCC 90028), *P. gingivalis* (ATCC 33277) and *F. nucleatum* (ATCC 25586) were all purchased as lyophilized stock (KWIK-STIK, VWR, Radnor, PA). Agar plates were also purchased from Sigma-Aldrich (St. Louis, MO). XTT Cell Viability Assay Kit was purchased from Biotium (Fremont, CA).

Citrate-capped silver nanoparticles (OD = 1) were purchased from Nanocomposix (San Diego, CA). Gold(III) chloride hydrate ($HAuCl_4 \cdot 3H_2O$, ≥99.9%), sodium citrate tribasic dihydrate (>99%), Trizma® base (>99.9%), Trizma® hydrochloride (>90%), DL-dithiothreitol (DTT, >99%), were purchased from Tokyo Chemical Industry Co., Ltd (TCI). Sodium chloride (NaCl, certified ACS), urea (certified ACS), and hydrochloric acid (certified ACS) were purchased from Fisher Chemical (Waltham, MA).

### II. Experimental Procedures

**Peptide synthesis.** Peptides were synthesized as prepared in previous work.[1] Briefly, an automated Eclipse™ peptide synthesizer (AAPPTec, Louisville, KY) was utilized for standard solid phase Fmoc synthesis on Rink-amide resin (0.55 mmol/g, 200 mg). Amino acids were coupled (C to N) under nitrogen protection with 0.2 M Fmoc-amino acid (5 equiv.) in 3 mL DMF, 0.2 M HBTU (5 equiv.) in 3 mL DMF, 0.4 M DIPEA (7.5 equiv.) in 3 mL DMF, and 20% (v/v) piperidine in 2 x 4 mL DMF for each coupling cycle. The resulting resin and peptide were then transferred to a syringe filter (Torviq Inc.) and washed with three rounds of DMF (5 mL each) and three rounds of DCM (4 mL each). It was then dried under vacuum. For acetylated peptides, the N-terminal was acetylated using the following recipe: 4 mL of DMF, 0.5 mL of Pyridine, and 0.5 mL of acetic anhydride. The solution was then subjected to light stirred for 30 minutes before being purged and washed with DMF and DCM as previously mentioned. The dried peptides were next cleaved from the resin using a 5 mL cocktail solution that consists of: 83% TFA, 5% H2O, 5% thioanisole, 5% phenol, and 2% µL EDDET. The incubated solution was gently rotated for 2 hours. The resin was then filtered and the filtrate containing the crude peptide was collected and precipitated using cold ethyl ether (20 mL, –20 ºC) and centrifuged three times (8,000 rpm, 3 minutes). Once the supernatant was removed, the precipitated pellets were dried and re-suspended using 10 mL of ACN/H2O mixtures wherein the percentage of ACN was controlled based on the solubility of the peptide.

**Peptide purification and characterization.** Peptide purification was conducted as done by Retout *et al*. [1] Once dry, the crude peptides were purified with a Shimadzu LC-40 HPLC system equipped with a LC-40D solvent delivery module, photodiode array detector SPD-M40, and degassing unit DGU-403. An injection of 2 mL was utilized with a Zorbax 300 BS, C18 column (5 µM, 9.4 × 250 mm) using an elution flow rate of 5 mL/ min over a 40-minute gradient from 10% to 95% acetonitrile in water (0.05% TFA). The peptide bond absorbance of 220 nm was monitored closely, and the elution was collected for characterization. Electrospray ionization mass spectrometry (ESI-MS) on the positive ion mode via the Micromass Quattro Ultima mass spectrometer provided by the Molecular MS Facility (MMSF) at UC San Diego was utilized with an MEOH/ H2O mixture (1:1, v/v) and an injection volume of 5 µL. Some compounds were characterized using matrix-assisted laser desorption/ionization-time of flight (MALDI-TOF) mass spectrometer using a linear positive mode. Here, a-Cyano-4-hydroxycinnamic acid (HCCA) was the matrix utilized at a ratio of 1:3 and 2 µL was placed and dried with a heat gun to be analyzed. Duplicates were used as recommended by the MMSF. Fractions containing the pure peptide as confirmed by electrospray ionization mass spectrometry (ESI-MS, positive ion mode) were lyophilized in a FreeZone Plus 2.5 freeze dry system (Labconco Corp., Kansas, MO) and aliquoted and stored in dry conditions at 2 ºC for further use.[2]

**Michaelis-Menten kinetics.** The peptide substrate was resynthesized with acetylated lysine residues and linked to cyanine-NHS ester dyes (Cy5.5 and Cy3) using the amine groups in the form of $NH_3^+$ and lysine at the N-terminal and the C-terminal, respectively: (V[aK][aK][aK]DVVDK). Acetylated lysine residues were employed to guide the dye to the lysine on the end terminal. Herein, 0.5 mg of the peptide dissolved in 369 µL of anhydrous DMSO with 31 µL of 1% (v/v) triethylamine (2.25 µmol, 1.5 equiv.) Next, Cy5.5 and Cy3 were added to the solution (3.5 µg, 4.50 µmol) and the solution was covered with aluminum foil and left to stir overnight at 300 rpm. The crude reaction was then dried under vacuum centrifugation using a Vacufuge Plus (Eppendorf, Hamburg) at 60 ºC until light-reflecting pellet was formed. The pellet was then resuspended in 25% ACN/ $H_2O$ (v/v) and separated using HPLC.

The heterodimer was diluted in 9.5 mM MES, 2.7 mM KCl, 140 mM NaCl (pH 5.5) buffer to reach a final [S] and distributed in 24 wells within a 96-well plate for duplicate measurements of each concentration. The enzyme ($[E]_0$ = 200 nM with respect to a final 100 µL volume) was then added to each well. The plate was incubated at 37 °C and the fluorescence intensity (684 nm for Cy5.5 and 570 nm for Cy3) was recorded over 12 h with 1 min intervals between each cycle. Measurements were performed in duplicates. The signal values at 30 min readout time were averaged and plotted against substrate concentrations; error bars represent the standard error of the means. The $\Delta PL = PL_{30\ min} - PL_{0\ min}$ was then correlated to product concentration using a standard curve: $\Delta PL_{Cy5.5}$ vs. [fully-digested FRET probe]. Data were then fitted to the following Michaelis-Menten equation: $v = \frac{V_{max}[S]}{K_{m+[S]}}$.

**Confocal Microscopy.** Mammalian cells were seeded at a density of $1\times10^6$ cells/ mL in 300 µL of their respective medium in 35 mm glass-bottom dishes (Cellvis, Lot No.: D35-14-1.5-N) and allowed to grow overnight. The cells were subsequently treated with 100 µM of the prodrug-peptides for 3 hours at 37 °C. Next, cells were washed three times with sterile PBS. Mammalian cells were stained at a final concentration of 1X CellBrite Fix Membrane Stain 640 (Biotium) at 37 °C for 15 minutes. Cells were then washed 3 times by PBS and fixed by 4% paraformaldehyde in PBS (Thermo Scientific) at room temperature for 20 min. Cells were imaged by a confocal microscope (Leica SP8 with lighting deconvolution) at an excitation/ emission wavelength of 638 nm/ 667 nm.

**Transmission Electron Microscopy.** TEM images were taken using the JEOL 1200 EX II operated at 80 kV. The TEM grids were prepared by drop casting 2 µL of each sample followed by air drying overnight. Cellular morphology was conducted using the same microscope. Here, 70nm thin sections of plastic were embedded with the fixed samples to visualize cellular organization.

**Mammalian cell culture.** HEK293T cells were a kind gift from Professor Liangfang Zhang's nanomedicine lab (UCSD NanoEngineering). Cells were cultured in complete Dulbecco's Modified Eagle Medium (supplemented with 10% Fetal Bovine Serum, 1% Penicillin-Streptomycin), at 37 °C and 5% CO2. Passages were conducted with 0.25% Trypsin EDTA before experiments.

**MRSA USA300 culture.** MRSA USA300 was a kind gift Professor Liangfang Zhang's nanomedicine lab (UCSD NanoEngineering) and was cultured on a tryptic soy broth (TSB) agar plate overnight at 37°C.[3] The bacteria were harvested by centrifugation at 5000 × g for 10 min before being used at $1 \times 10^6$ cells/ mL (OD600 = 1.0, logarithmic growth phase) in a 96-well plate for viability assay experimentation.

**Limit of Detection.** The limit of detection (LoD) was calculated using the limit of blank (LoB) as demonstrated by Armbruster et al.[4] The LoB uses a blank measurement to define the highest signal generated from the sample with no analyte. LoB was calculated using the mean ($mean_{blank}$) and standard deviation ($SD_{blank}$) of a blank sample:

$LoB = mean_{blank} + 1.645\ (SD_{blank})$

Based on this, the LoD is defined as the lowest analyte concentration that can be differentiated from the LoB. Here, the LoD represents an analyte concentration at which 95% of measured samples are readily differentiated from the LoB while the remaining 5% can contain no analyte:

$LoD = LoB + 1.645\ (SD_{low\ concentration\ sample})$

III. Supplementary Data

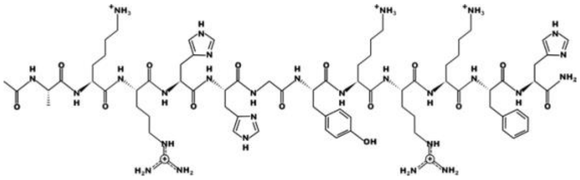

**P-113**: AKRHHGYKRKFH (1604.8891 g/mol)

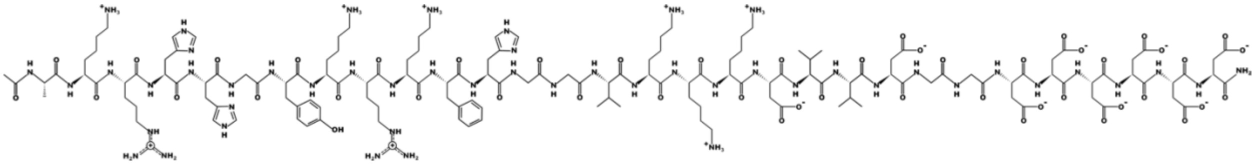

**S1**: AKRHHGYKRKFHGGVKKKDVVDGGDDDDDD (3434.6778 g/mol)

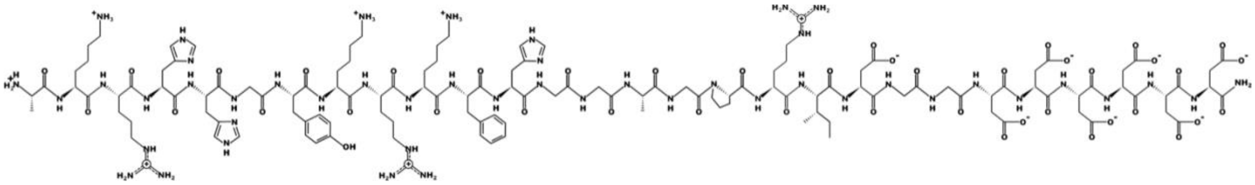

**G1**: AKRHHGYKRKFHGGAGPRIDGGDDDDDD (3090.4474 g/mol)

**Figure S1. Structure of P-113 and SAP9 and RgpB- cleavable prodrugs, S1 and G1, respectively (top to bottom).**

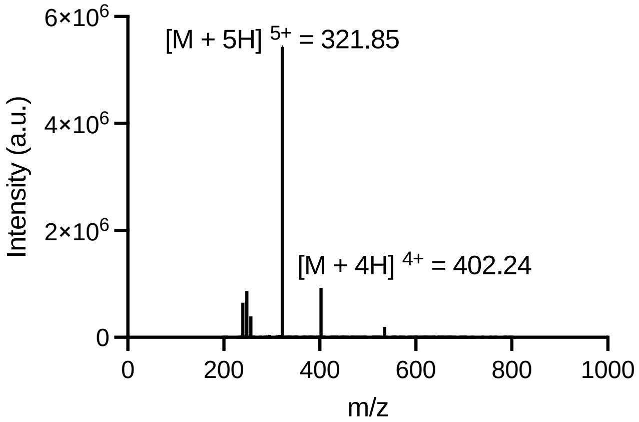


**Figure S2. P-113 characterization.** ESI-MS with theoretical mass: 1604.8891 Da, $[M + 5H]^{5+}$ = 321.98 m/z; $[M + 4H]^{4+}$ = 402.22 m/z.

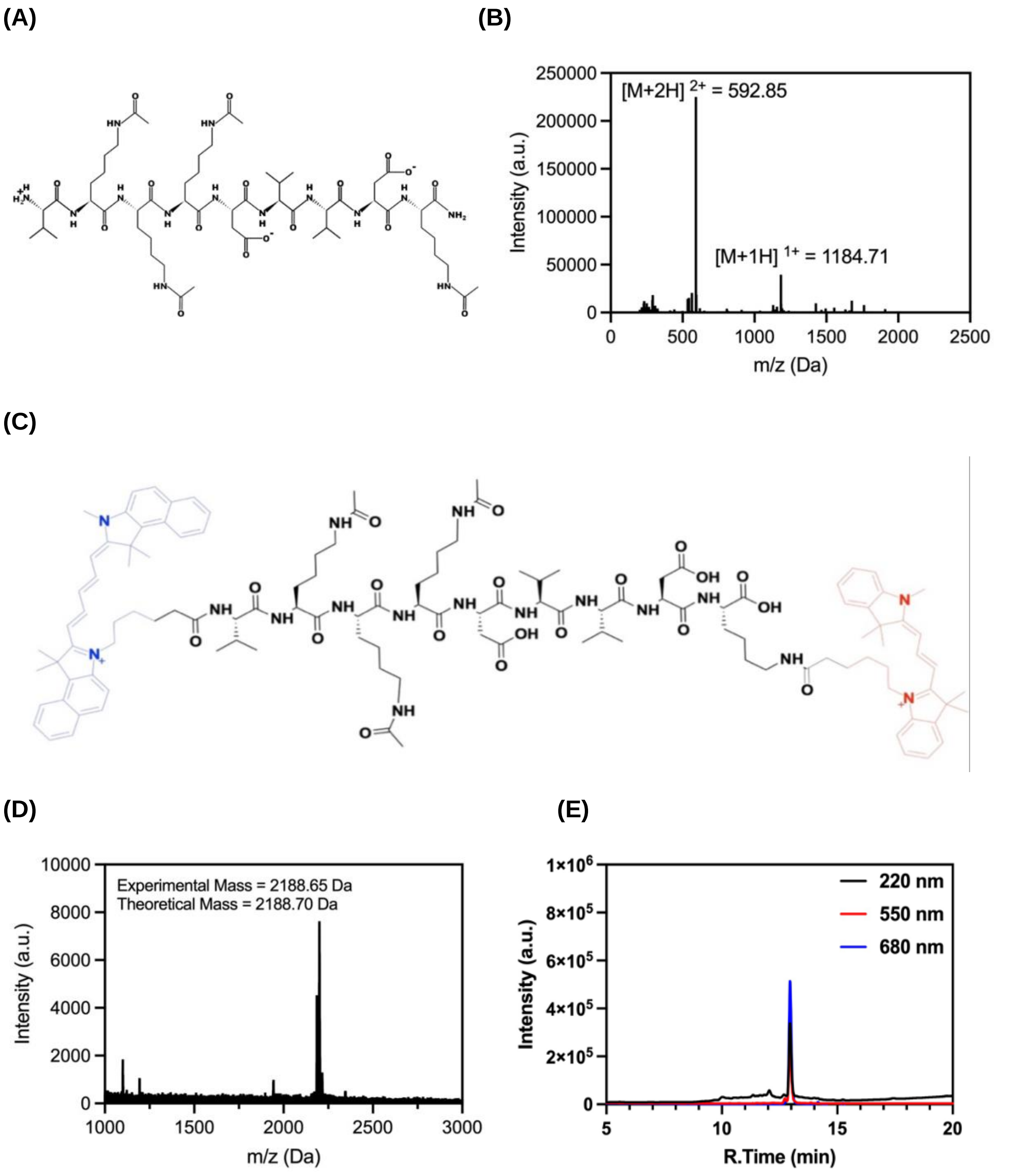


**Figure S3. SAP9 cleavable substrate FRET probe characterization. (A)** SAP9 cleavable substrate peptide structure: Val- aLys- aLys- aLys- Asp- Val -Val -Lys with theoretical mass 1182.6953 Da. **(B)** ESI-MS presenting two clear peaks at $[M+2H]^{2+}$ = 592.34765 and $[M+1H]^{1+}$ = 1183.6953. **(C)** Structure of SAP9 cleavable probe with conjugated NHS-dyes on either terminal with theoretical mass 2188.8039 Da. **(D)** MALDI-TOF spectra **(E)** HPLC spectra monitoring at peptide bond wavelength, 220 nm, and cyanine dye wavelengths, 550 nm and 680 nm confirming successful conjugation.

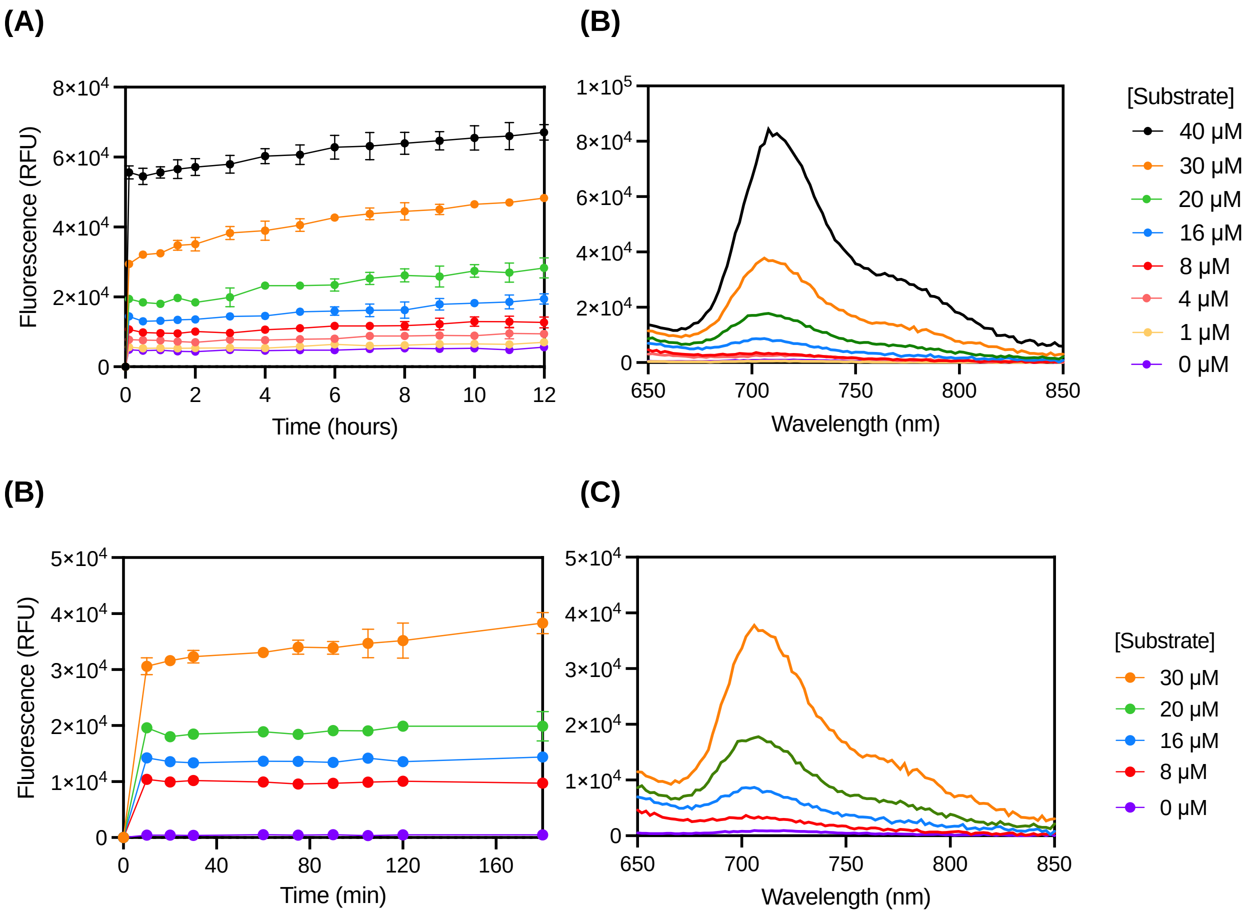


**Figure S4. SAP9 cleavable substrate FRET probe fluorescence. (A)** Kinetic data collected over 12-hour period. **(B)** Total corresponding fluorescence emission spectra (Ex.620 nm) after 3 hours incubation at 37°C at increasing concentration of substrate. **(C)** Extracted data for 3-hour incubation demonstrating maximum fluorescence reached. **(D)** Zoomed in (0 – 30 μM) corresponding fluorescence emission spectra (Ex.620 nm) after 3 hours incubation at 37°C at increasing concentration of substrate.

(A) (B)

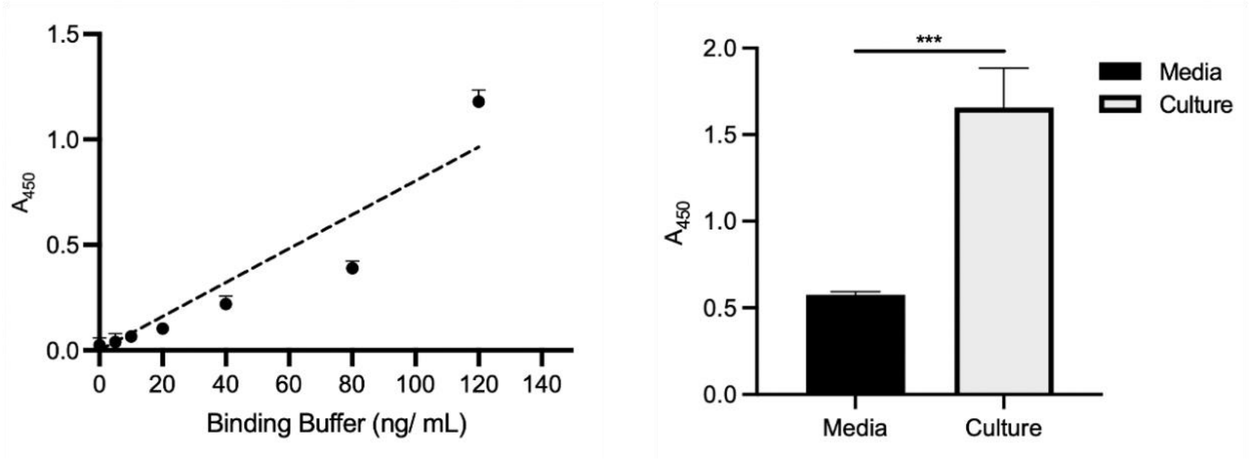


**Figure S5. ELISA detection of SAP in *C. albicans* culture. (A)** Standard curve for SAP detected via commercially available ELISA where m = $6.8 \times 10^{-3} \pm 0.1 \times 10^{-4}$. **(B)** C.a. culture (OD = 0.5) contained significantly more SAP than media alone. [SAP] from cells = 243 ± 33 ng/mL. Error bars represent the standard error for the mean for n = 6. Asterisks denote values from a one-tailed t-test (***$p < 0.0001$).

**(A)** **(B)**

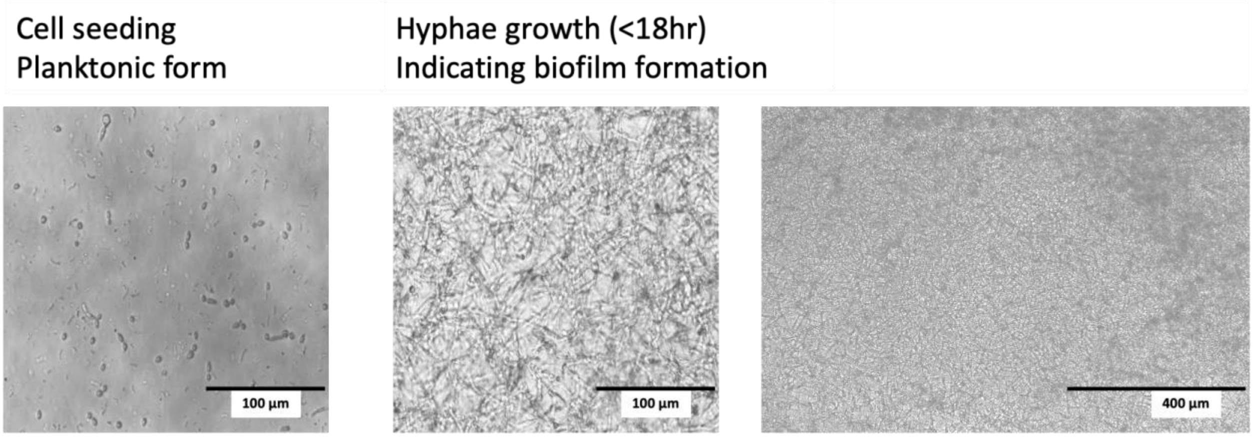


**Figure S6. *C. albicans* growth transmission microscopy. (A)** *C. albicans* cell seeding planktonic form (100 µm) and **(B)** hyphae growth (<18 hr.) indicating biofilm formation after 18 hours at 30°C (100 µm) and expanded to 400 µm.

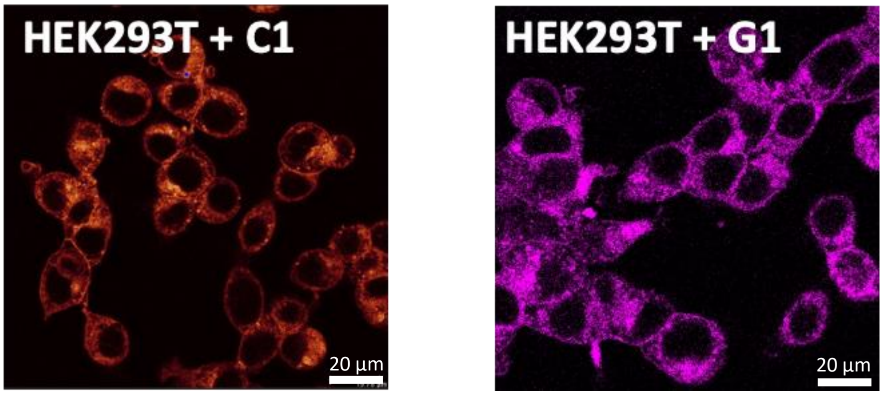


**Figure S7. Confocal microscopy of mammalian cells.** HEK293T demonstrating intact cell membrane after treatment with prodrugs S1 and G1 for 3 hours at 37 °C. Cells were stained at a final concentration of 1X at 37 °C for 15 minutes with CellBrite Fix Membrane Stain 640, fixed with 4% paraformaldehyde, and imaged with a Leica SP8 with lighting deconvolution at an excitation/ emission wavelength of 638 nm/ 667 nm.

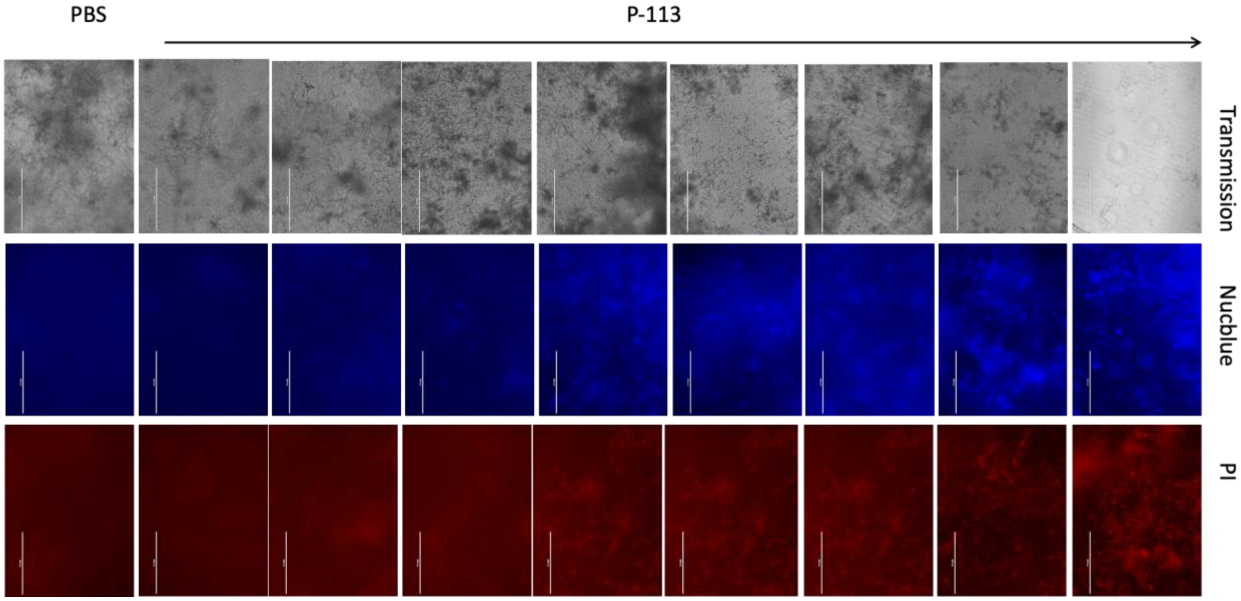


**Figure S8. Transmission, NucBlue, and propidium iodide microscopy for *C. albicans*.** 100 µm for *C. albicans* treated cells using primary treatment, S1. Microscopy demonstrates the effects of S1 on the biofilm formation. Compared to PBS, it is evident that increasing concentration results in the breakdown of the *C. albicans* hyphal structures and reversion to the planktonic form. NucBlue and propidium iodide staining were also used. NucBlue staining is a commonly used fluorescent dye that specifically binds to DNA, allowing for the visualization and quantification of nuclei within cells. The increased fluorescence of NucBlue associated with the increase in peptide concentration indicates an increase in the amount of DNA released as a result of cell death. Propidium iodide is another fluorescent dye that selectively binds to DNA, allowing for the detection of cells with compromised cell membranes. Thus, the increase in fluorescence signal indicates an increase in the number of cells with damaged or permeabilized membranes.

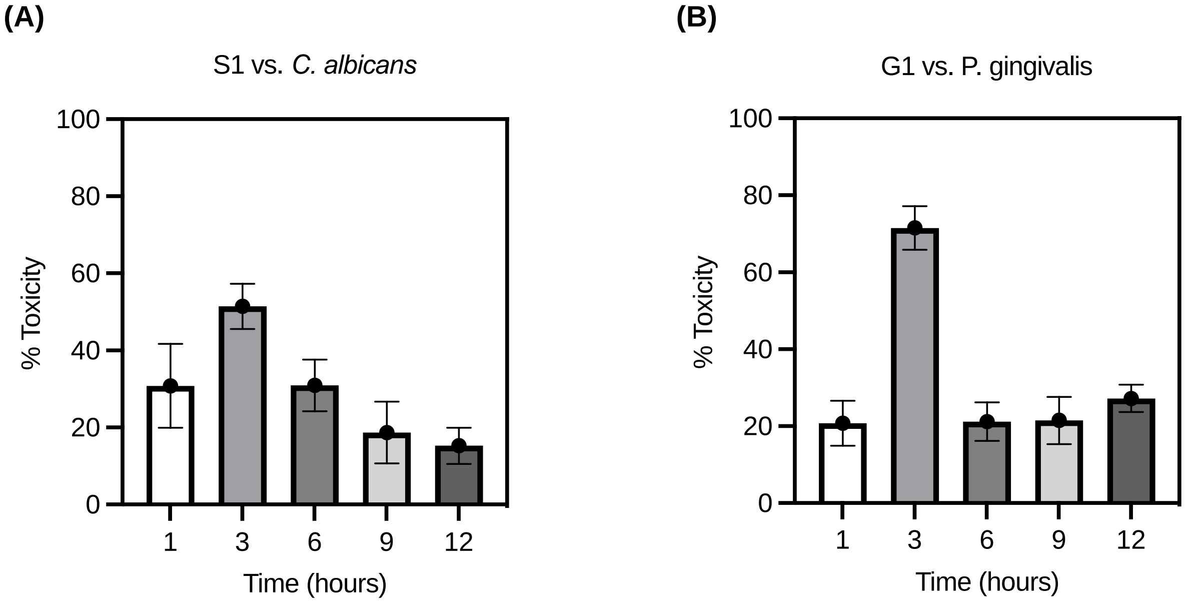


**Figure S9. Time dependent toxicity assessment for S1 and G1. (A)** XTT viability assay for *C. albicans* cells incubated with 10 µM of S1 in MES buffer over time. Here, an increased toxicity is observed at 3 hours. **(B)** XTT viability assay for *P. gingivalis* cells incubated with 10 µM of G1 in Tris buffer over time. Again, an increased toxicity is observed at 3 hours. Cells were washed with PBS to prevent continued antimicrobial activity past the desired time.

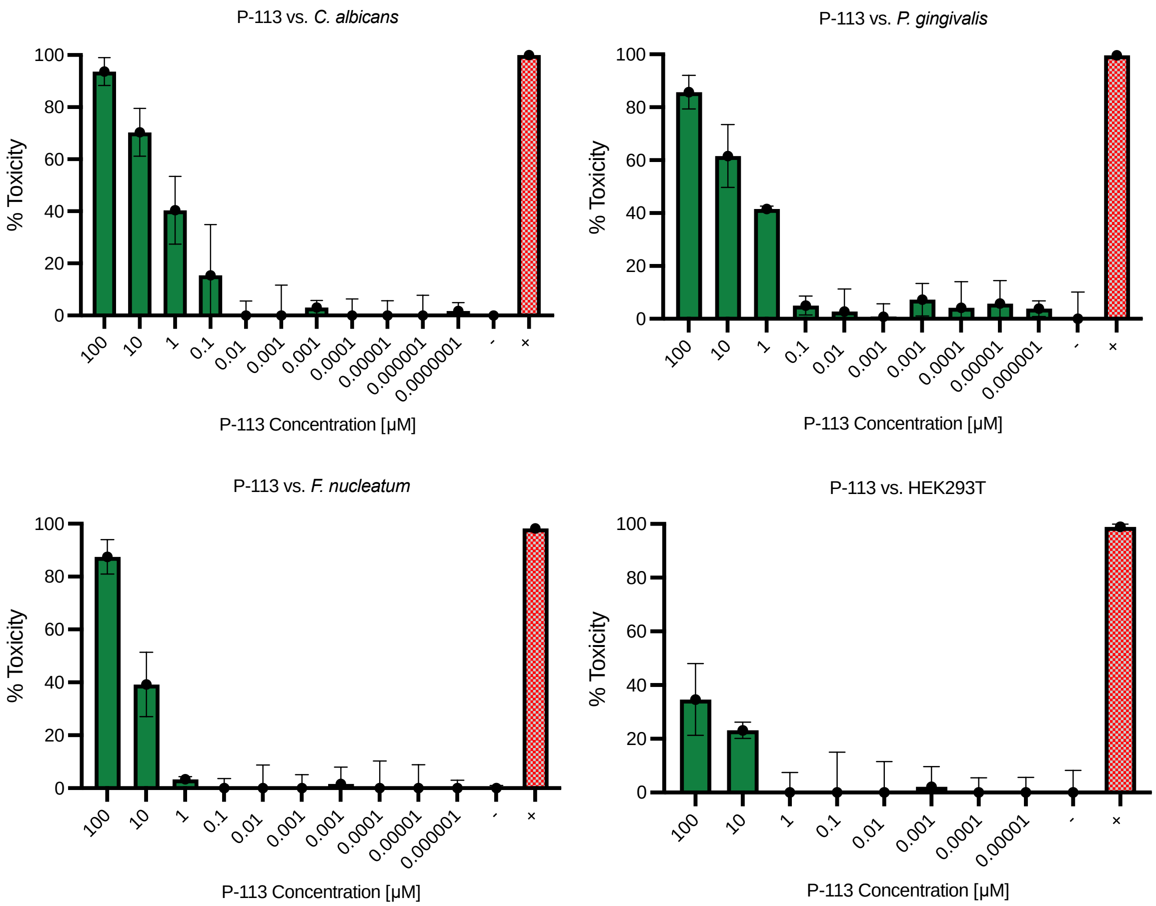


**Figure S10. Toxicity assessment for varying concentrations of P-113, positive control, treatment along four cell lines. (A)** *C. albicans* with negative control (-) MES buffer and positive control (+) Fluconazole. **(B)** *P. gingivalis* with negative control (-) Tris buffer and positive control (+) Penicillin-streptomycin. **(C)** *F. nucleatum* with negative control (-) Tris buffer and positive control (+) Penicillin-streptomycin. **(D)** HEK293T with negative control (-) PBS and positive control (+) 10% bleach.

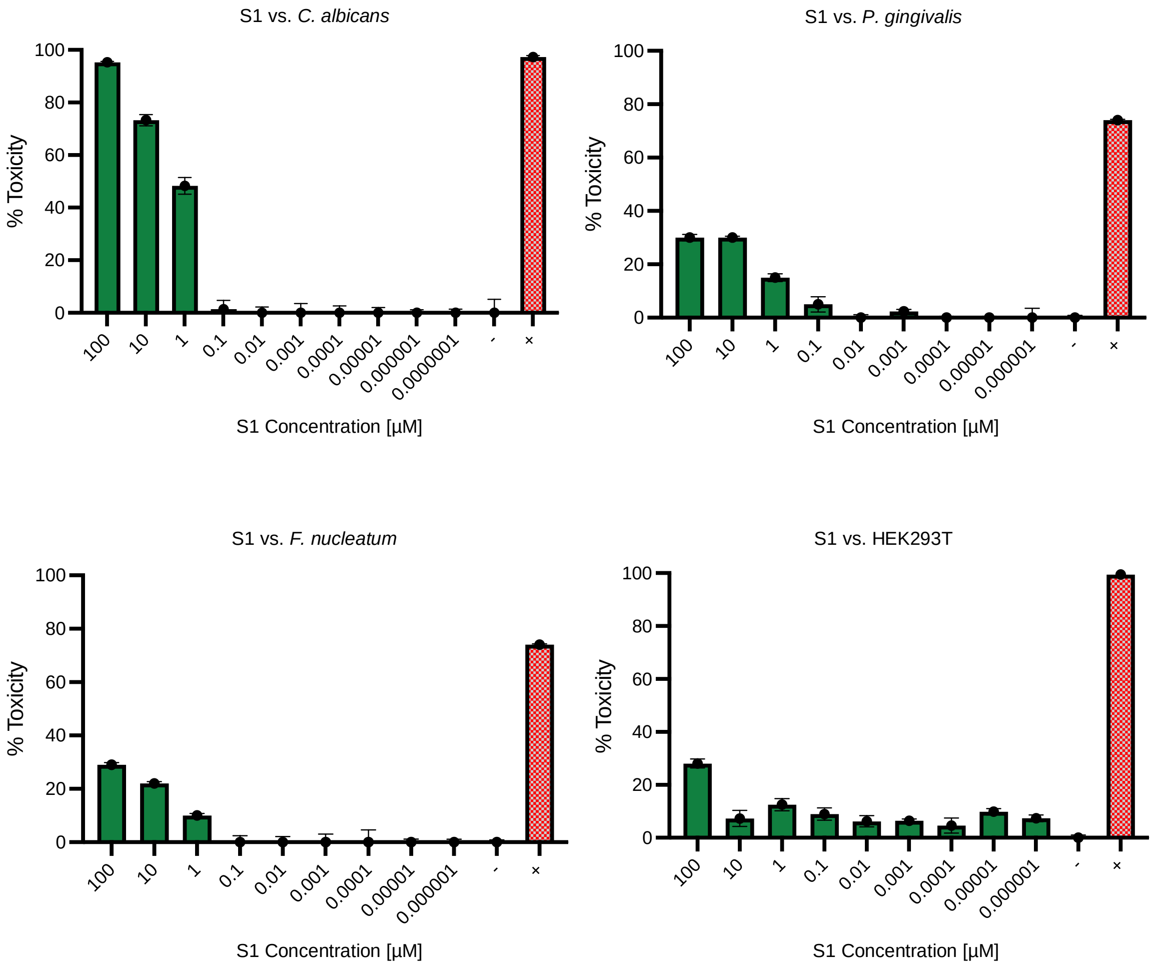


**Figure S11. Toxicity assessment for varying concentrations of S1, C. albicans prodrug, treatment, along four cell lines. (A)** *C. albicans* with negative control (-) MES buffer and positive control (+) Fluconazole*.* **(B)** *P. gingivalis* with negative control (-) Tris buffer and positive control (+) Penicillin-streptomycin*.* **(C)** *F. nucleatum* with negative control (-) Tris buffer and positive control (+) Penicillin-streptomycin. **(D)** HEK293T with negative control (-) PBS and positive control (+) 10% bleach.

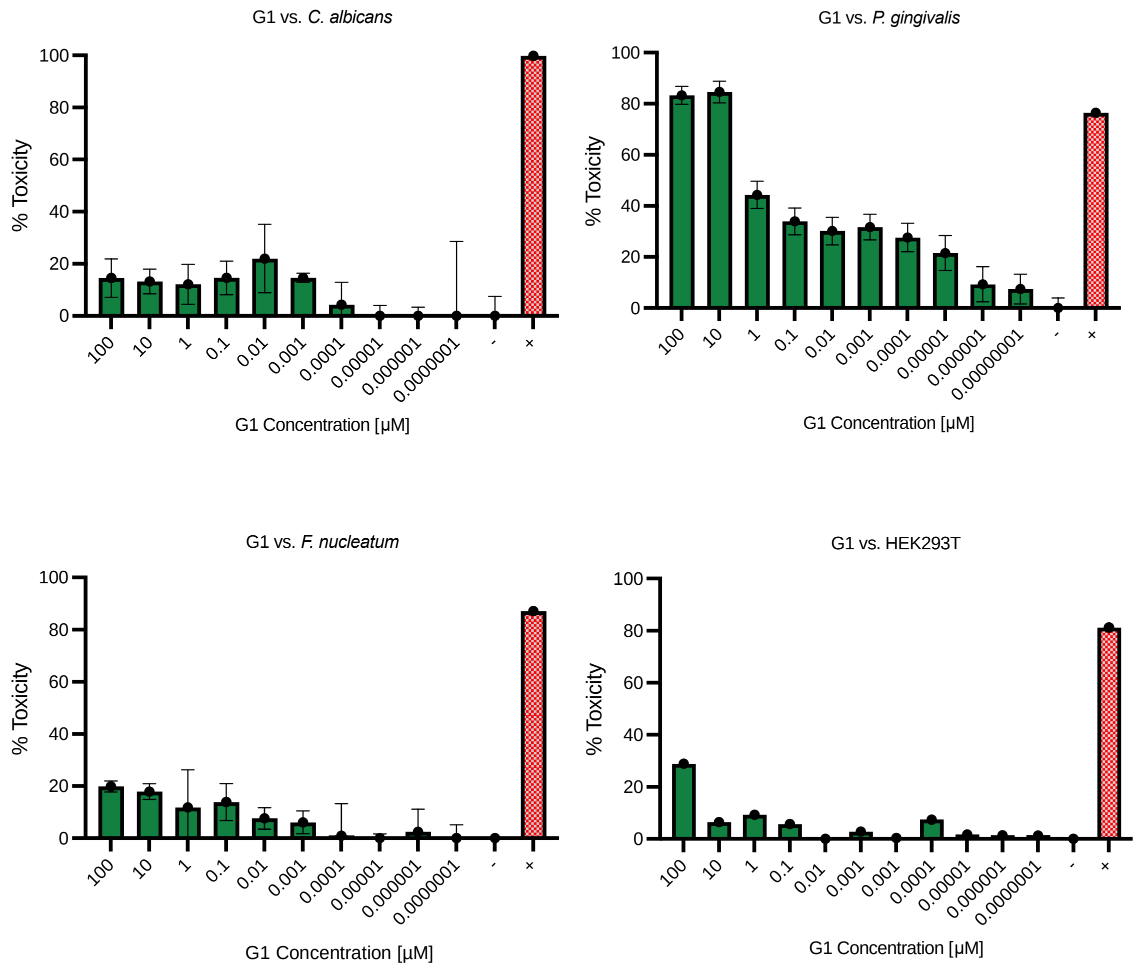


**Figure S12. Toxicity assessment for varying concentrations of G1, P. gingivalis prodrug, treatment, along four cell lines. (A)** *C. albicans* with negative control (-) MES buffer and positive control (+) Fluconazole. **(B)** *P. gingivalis* with negative control (-) Tris buffer and positive control (+) Penicillin-streptomycin. **(C)** *F. nucleatum* with negative control (-) Tris buffer and positive control (+) Penicillin-streptomycin. **(D)** HEK293T with negative control (-) PBS and positive control (+) 10% bleach.

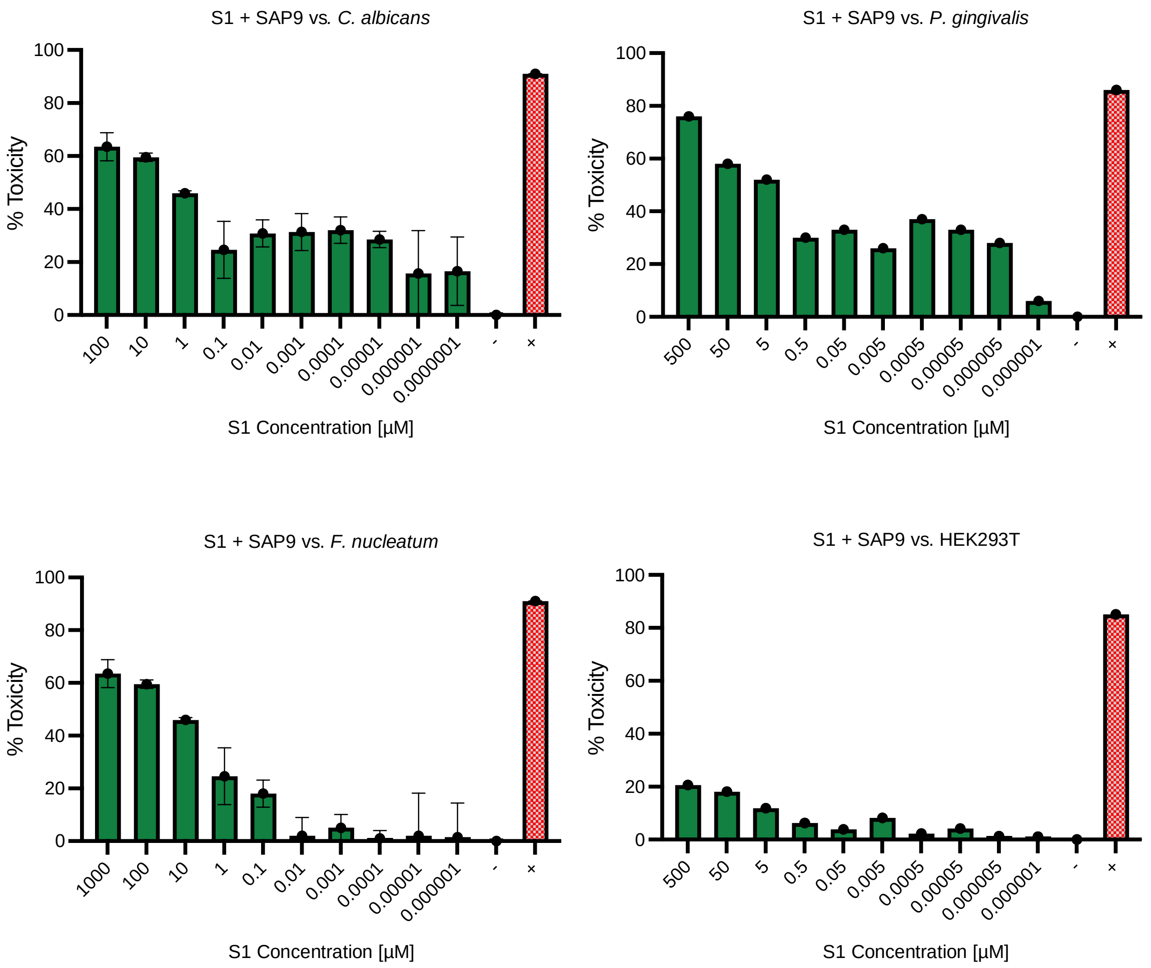


**Figure S13. Toxicity assessment for varying concentrations of S1 + SAP9, C. albicans prodrug pre-cleaved by 200 nM recombinant protease in vitro, treatment, along four cell lines. (A)** *C. albicans* with negative control (-) MES buffer and positive control (+) Fluconazole. **(B)** *P. gingivalis* with negative control (-) Tris buffer and positive control (+) Penicillin-streptomycin. **(C)** *F. nucleatum* with negative control (-) Tris buffer and positive control (+) Penicillin-streptomycin. **(D)** HEK293T with negative control (-) PBS and positive control (+) 10% bleach.

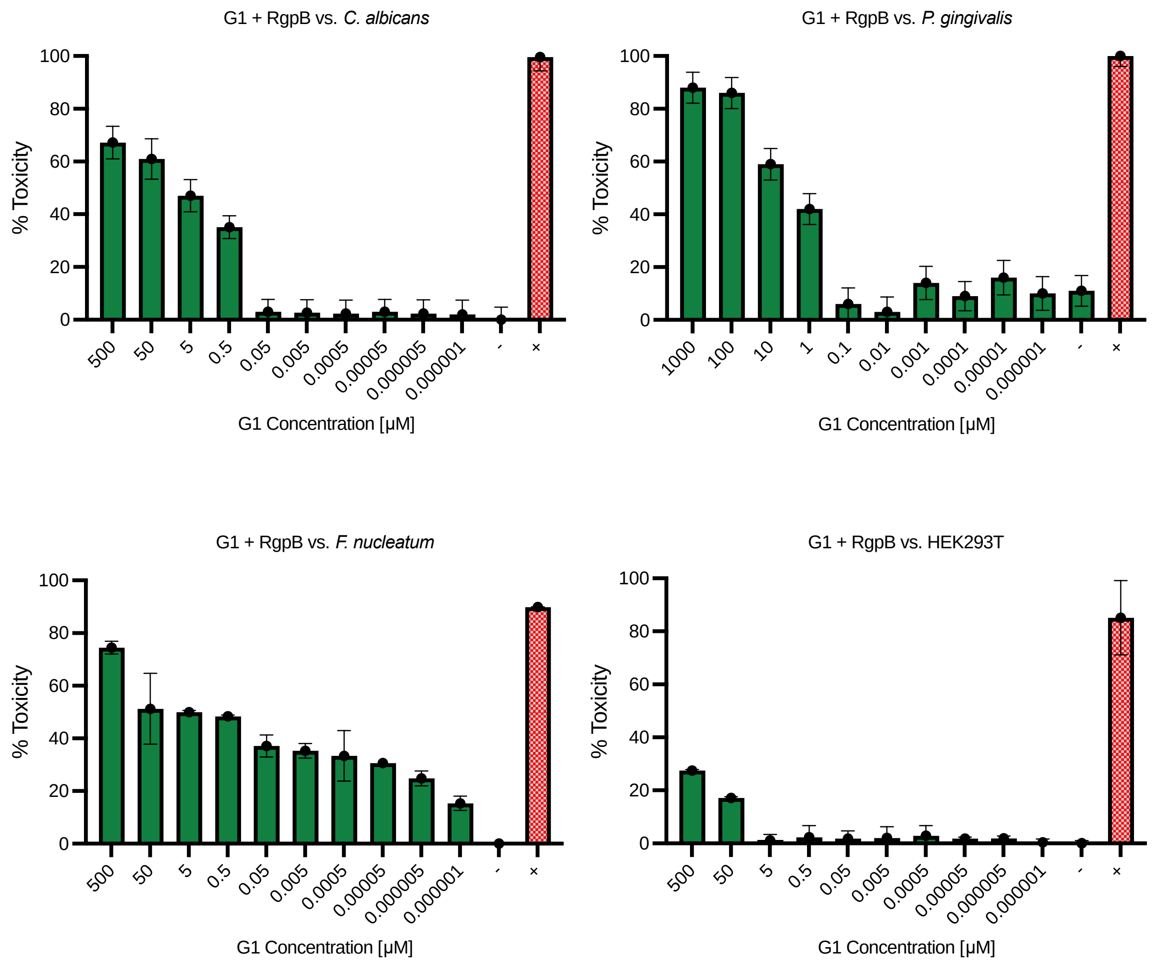


**Figure S14. Toxicity assessment for varying concentrations of G1 + RgpB, P. gingivalis prodrug pre-cleaved by 200 nM recombinant protease in vitro, treatment, along four cell lines. (A)** *C. albicans* with negative control (-) MES buffer and positive control (+) Fluconazole. **(B)** *P. gingivalis* with negative control (-) Tris buffer and positive control (+) Penicillin-streptomycin. **(C)** *F. nucleatum* with negative control (-) Tris buffer and positive control (+) Penicillin-streptomycin. **(D)** HEK293T with negative control (-) PBS and positive control (+) 10% bleach.

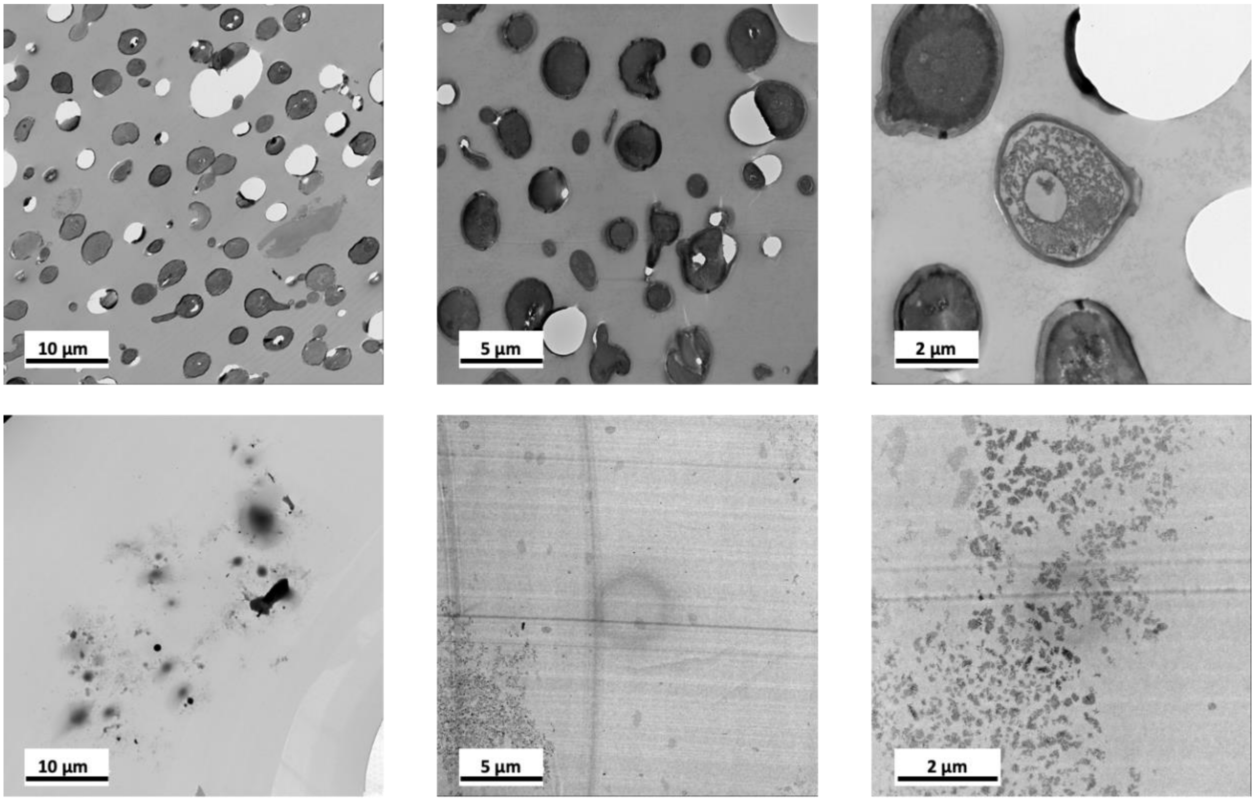


**Figure S15. C. albicans TEM images of cellular morphology at increased magnification.** Top: Intact cells alone with intact membranes. Bottom: Cells with damaged membranes after incubation with S1 for 3 hours at 37 °C (increasing magnification left to right).

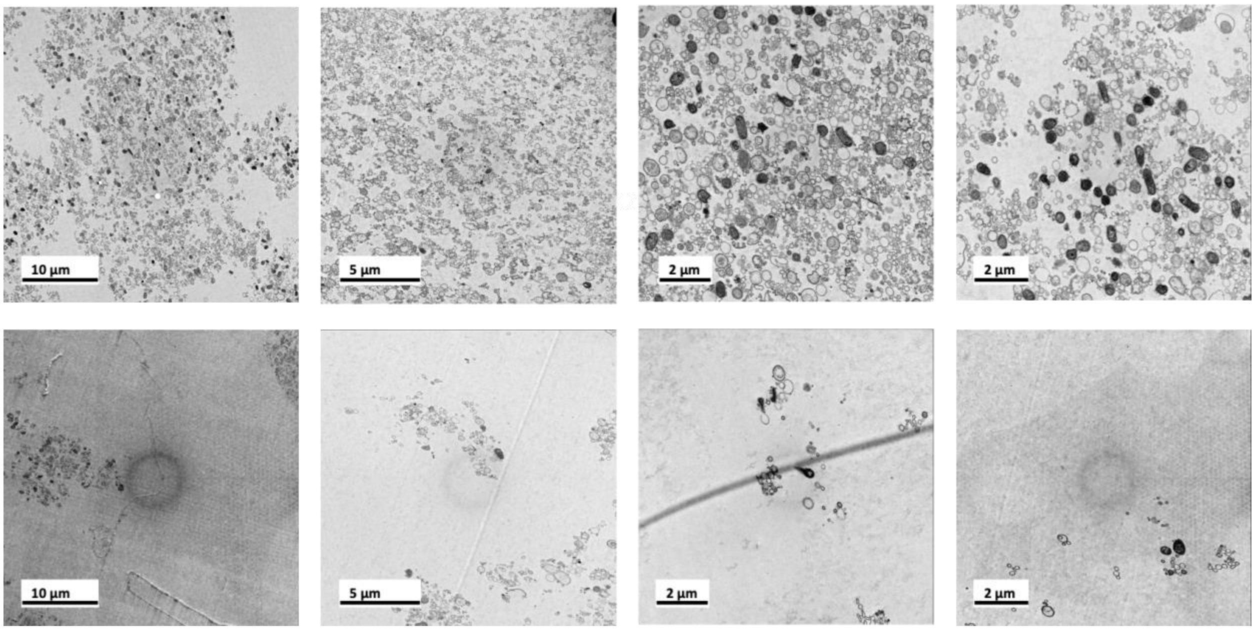


**Figure S16. P. gingivalis TEM images at increased magnification.** Top: Intact cells alone at high density. Bottom: Deconstructed cells after incubation with G1 for 3 hours at 37 °C (increasing magnification left to right).

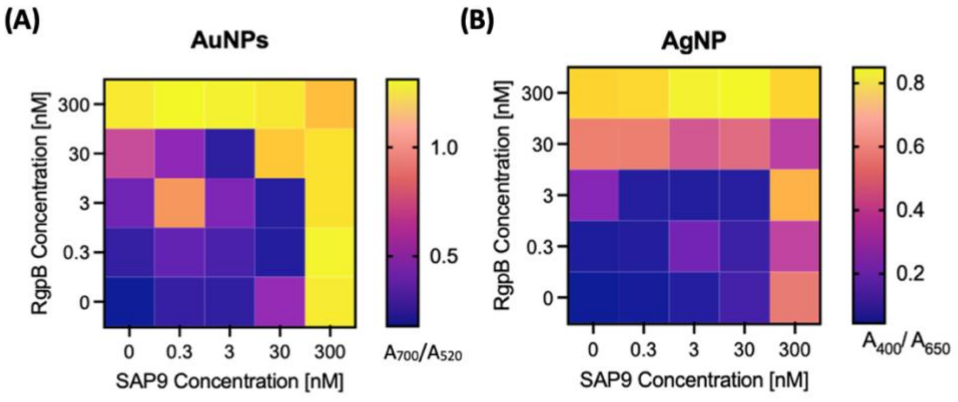


**Figure S17. Heat map for varying protease concentration in LOD analysis. (A)** Gold nanoparticle and **(B)** silver nanoparticle limit of detection for SAP9/ RgpB at increasing concentration from 0 – 300 nM. At the highest concentration of both proteases, nanoparticle assembly is increased as presented by the increase in absorbance ratio and by-eye color change from red to blue.

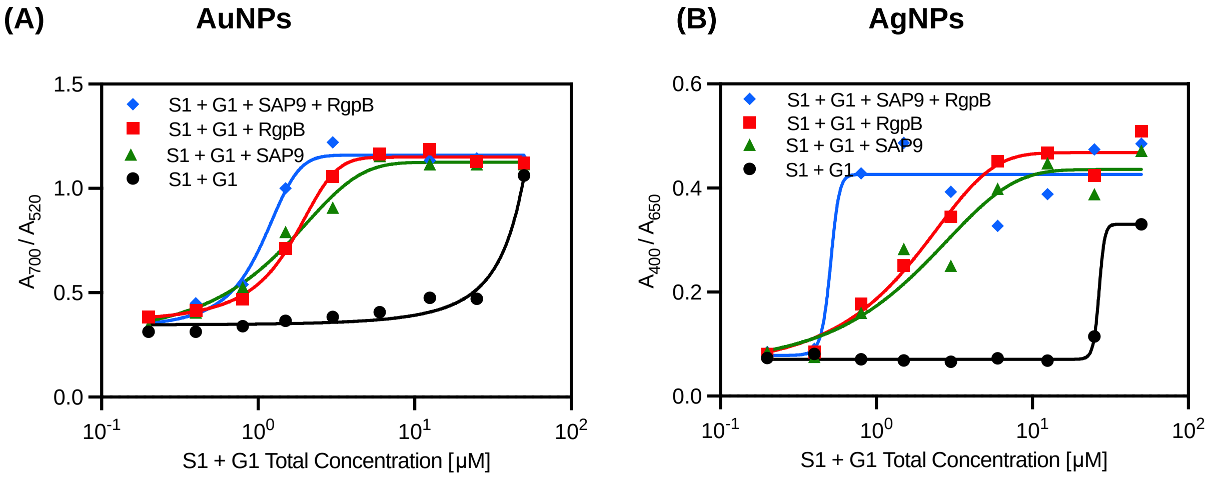


**Figure S18. Colorimetric detection titration curves for combined S1 and G1. (A)** Optical absorption of AuNPs and **(B)** AgNPs when incubated with increased concentrations of S1 and G1 co-incubated (black) and variations of fragmented peptides. With both enzymes (blue), the system is more sensitive as represented by the more rapid increase in absorbance. Single-enzyme incubation with both peptides returns a steady increase with a clear difference to the no-enzyme conditions.

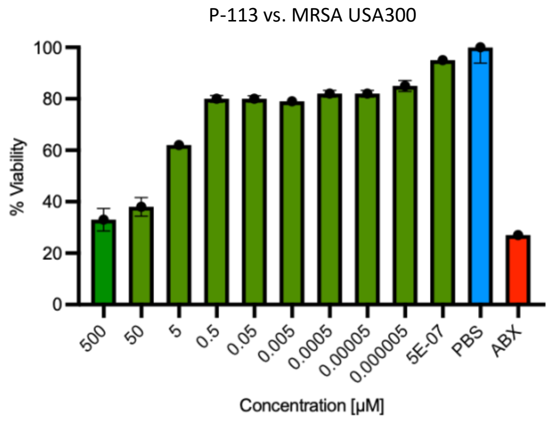


**Figure S19. Gram-positive bacteria toxicity assay of P-113 against MRSA USA300.** MRSA USA300 with negative control (-) PBS and positive control (+) Penicillin-streptomycin. Although gram-positive bacteria are more susceptible to P-113 AMP mechanism, MRSA strains are highly resistant, yielding a higher MIC for P-113 compared to the investigated gram-negative oral microorganisms.[5, 6]

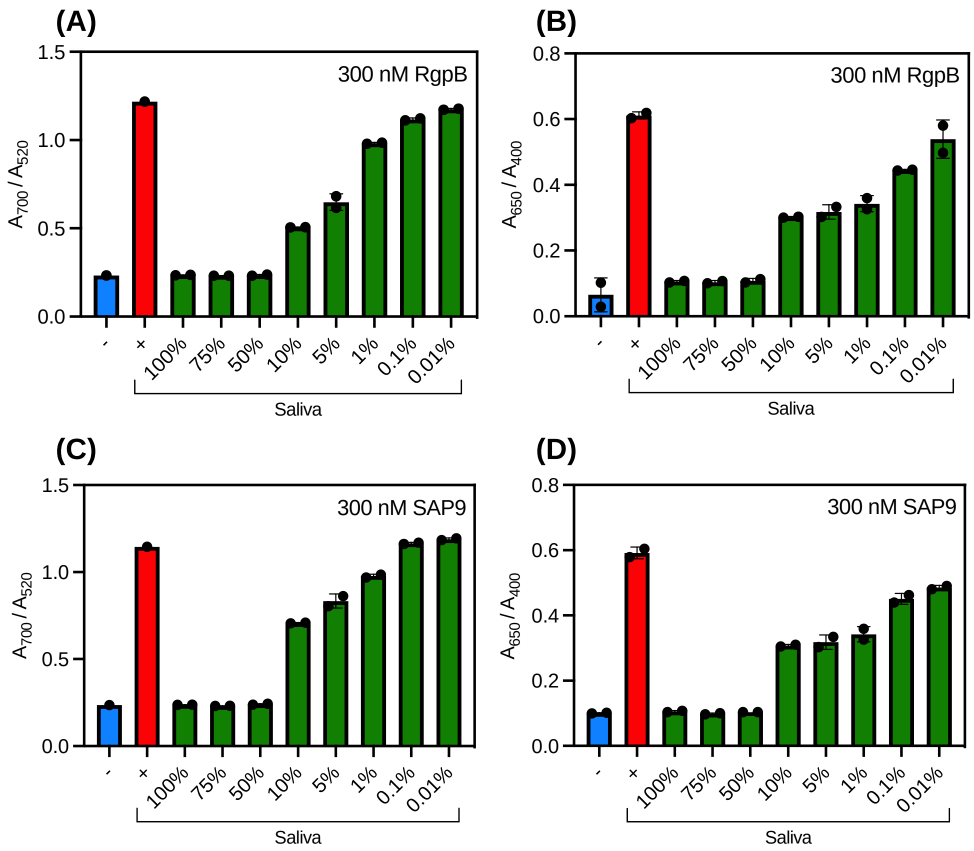


**Figure S20. Protease-induced plasmonic nanoparticle assembly in saliva. (A)** Optical absorption of AuNPs and **(B)** AgNPs when incubated with increased concentrations of saliva spiked with 300 nM RgpB in activity buffer at constant G1 concentration (1:1000, E:P). **(C)** Optical absorption of AuNPs and **(D)** AgNPs when incubated with increased concentrations of saliva spiked with 300 nM SAP9 in activity buffer at constant S1 concentration (1:1000, E:P).

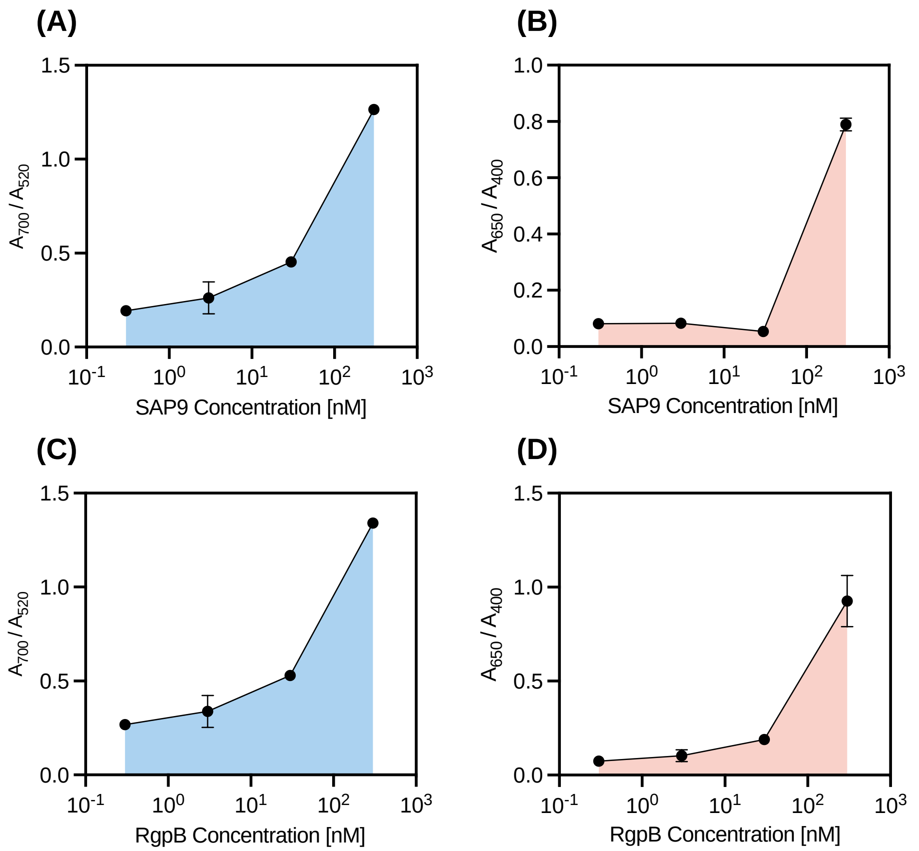


**Figure S21. Limit of detection of plasmonic nanoparticle assembly in saliva. (A)** Limit of detection (LOD) for AuNP and **(B)** AgNP detection system of S1 using an increasing concentration of SAP9. **(C)** Limit of detection (LOD) for AuNP and **(B)** AgNP detection system of G1 using an increasing concentration of RgpB. LOD study shows an increase in the limit for both proteases in complex media.

### V. Author Contributions

L.A. proposed the system, designed, and conducted experiments, led data curation and analysis, and wrote the manuscript. M.R. helped design the system, conduct experiments, and analyze data. J.V.J. led project administration, and conceived, and supervised the work. All authors edited the manuscript.
